# The Condensin II Complex Regulates the Expression of Essential Gene Expression Programs During Erythropoiesis

**DOI:** 10.1101/2024.11.05.621908

**Authors:** Deanna Abid, Kristin Murphy, Zachary Murphy, Nabil Rahman, Michael Getman, Laurie Steiner

## Abstract

Erythropoiesis is characterized by dramatic changes in gene expression in the context of a cell that is rapidly proliferating while simultaneously condensing its nucleus in anticipation of enucleation. The mechanisms that maintain high level expression of erythroid genes and promote nuclear condensation remain poorly understood. Condensin II is a ring-like complex that promotes mitotic chromatin condensation and has roles in regulating interphase chromatin architecture and gene expression. We interrogated the role of Condensin II in erythropoiesis using an erythroid-specific deletion of the Condensin II subunit, Ncaph2. Ncaph2 loss resulted in severe embryonic anemia with lethality at embryonic day 13.5. Ncaph2 mutant erythroid cells had dysregulated maturation and disrupted cell cycle progression, but surprisingly Ncaph2 was dispensable for nuclear condensation. Genomic studies revealed that Ncaph2 occupied the promoter of key erythroid and cell cycle genes that were downregulated following Ncaph2 loss. Together, our results demonstrate an essential role for Ncaph2 in the gene expression programs that regulate cell cycle progression and erythroid differentiation, and identify a key role for the Condensin II complex in the regulation of a lineage-specific differentiation program.

**Summary Statement:** The Condensin II complex regulates cell cycle progression and erythroid differentiation, but is dispensable for nuclear condensation during terminal erythroid maturation.

## Introduction

The Condensin II complex is a multi-subunit, ring-like structure that plays a key role in chromosome compaction and segregation during mitosis. Condensin II is composed of two Structural Maintenance of Chromosomes (SMC) subunits, Smc2 and Smc4, along with Ncapd3 (non-SMC Condensin II subunit D3), Ncapg2 (non-SMC Condensin II subunit G2), and the kleisin protein Ncaph2 (non-SMC Condensin II subunit H2). The Condensin II complex has a well described role in cell cycle progression, contributing to chromosome condensation and segregation during mitosis. (Hirota et al., 2004, Ono et al., 2004, Ono et al., 2017) In contrast to the related Condensin I complex, which is excluded from the nucleus until nuclear envelope breakdown during mitosis, the Condensin II complex is present in the cell throughout interphase, with several studies implicating it in the regulation of interphase chromatin architecture (Hirota et al., 2004, Ono et al., 2004, Hoencamp et al., 2021, Nishide and Hirano, 2014). Studies on the role of Condensin II in the regulation of gene expression have been contradictory, with some associating Condensin II with both gene activation and repression, and others finding minimal changes in gene expression following Condensin II disruption. (Hoencamp et al., 2021, Woodward et al., 2016) The role of the Condensin II complex in gene regulation remains incompletely understood, and few studies have assessed the function of the Condensin II complex *in vivo*. (Nishide and Hirano, 2014, Woodward et al., 2016, Macdonald et al., 2022)

Erythropoiesis is the dynamic process of generating enucleated red blood cells from hematopoietic stem and progenitor cells. During terminal maturation of erythroid cells, they must condense their nuclei to 1/10^th^ of their original volume in preparation for enucleation.(Ji et al., 2011) In the context of these dramatic changes in nuclear architecture, erythroid precursors must silence the majority of the non-erythroid genome while maintaining high-level expression of globins and other transcripts essential for the erythroid differentiation program. The mechanisms that coordinate gene expression, cell cycle progression, and nuclear condensation remain incompletely understood. We hypothesized that the Condensin II complex may play an important role in regulating these processes. The subunits of the Condensin II complex, including Ncaph2, are highly expressed in maturing erythroid cells compared to other hematopoietic cell types.(Bagger et al., 2019) Further, overexpression of the Condensin II subunit NCAPG2 accelerates erythroid differentiation in Murine Erythroleukemia (MEL) cells.(Xu et al., 2006) Together these data suggest a role for the Condensin II complex in regulating erythropoiesis.

We have previously shown that loss of the histone methyltransferase, Kmt5A (Setd8, pr-Set7) results in severe embryonic (primitive erythroid) anemia due to loss of chromatin condensation and disrupted erythroid maturation.(Malik et al., 2017, Myers et al., 2020) Kmt5A-mediated H4K20 mono-methylation (H4K20me1) is an important mediator of higher order chromatin condensation. (Beck et al., 2012, Oda et al., 2009, Evertts et al., 2013) Further, studies in cell lines suggest that H4K20me1 can recruit members of the Condensin II complex, implying that this interaction may be important for mitotic chromatin condensation.(Liu et al., 2010) We sought to understand the function of the Condensin II complex and its potential interaction with H4K20me1 during erythropoiesis. To interrogate the role of Condensin II in a primary model, we crossed mice expressing cre-recombinase under the direction of the endogenous erythropoietin receptor promoter [EporCre;(Heinrich et al., 2004)] with mice containing a floxed allele of the Condensin II subunit Ncaph2, which is essential for the assembly of the complex (Hoencamp et al., 2021). Embryos homozygous for EpoR-specific excision of *Ncaph2* suffered from severe anemia with lethality by embryonic day (E) 13.5. In stark contrast to Kmt5a mutant erythroblasts, nuclear condensation was maintained in Ncaph2 mutant cells, although erythroid maturation and gene expression were both disrupted. Genomic studies demonstrated that Ncaph2 occupied the transcription start sites of highly expressed erythroid genes as well as genes essential for cell cycle progression. Together, these studies identify a role for the Condensin II complex in the regulation of a lineage-specific differentiation program.

## Results

### Ncaph2 is Essential for Erythropoiesis

As with the other subunits of Condensin II, Ncaph2 is highly expressed in terminally maturing erythroblasts (Fig S1). To determine the role of the Condensin II complex in erythropoiesis we crossed mice harboring floxed alleles of Ncaph2 (*Ncaph2* fl/fl) with mice expressing Cre recombinase under the direction of the endogenous erythropoietin Receptor promotor (EpoRCre; (Heinrich et al., 2004)]. The resulting embryos with conditional erythroid deletion of *Ncaph2* (*Ncaph2* 11/11) had mild pallor at E11.5 (embryonic day 11.5), and dramatic pallor by E13.5 (Fig 1A). Peripheral blood counts revealed a severe defect in the number of primitive erythroblasts in *Ncaph2* 11/11 embryos (Fig 1B) and flow cytometric analyses with antibodies targeting CD71 and Ter119 suggested delayed maturation (Fig 1C, S1). Primitive erythroid cells mature semi-synchronously in circulation, characterized by a progressive decrease in cell and nuclear size and accumulation of hemoglobin (Palis et al., 2010). Despite their immunophenotypic delay in maturation, the *Ncaph2* 11/11 cells had an acidophilic appearance consistent with hemoglobin accumulation, and quantitative PCR demonstrated similar levels of globin gene expression in all genotypes (Figs 1D, E). As expected, the control cells were very uniform in appearance. In contrast, the Ncaph2 mutant primitive erythroid cells had more variation in cell size (Figs 1D,F), and measurement of cell size by imaging flow cytometry revealed a modest increase in average cell size in the *Ncaph2* 11/11 erythroblasts compared to littermate controls (Fig 1F). Together, these data suggest that Ncaph2 loss dysregulates but does not block primitive erythroid maturation. On the other hand, the fetal liver appears to be absent in E12.5 and E13.5 *Ncaph2* 11/11 embryos, suggesting a severe defect in definitive erythropoiesis that results in a more profound anemia (Fig 1A).

**Figure 1.**
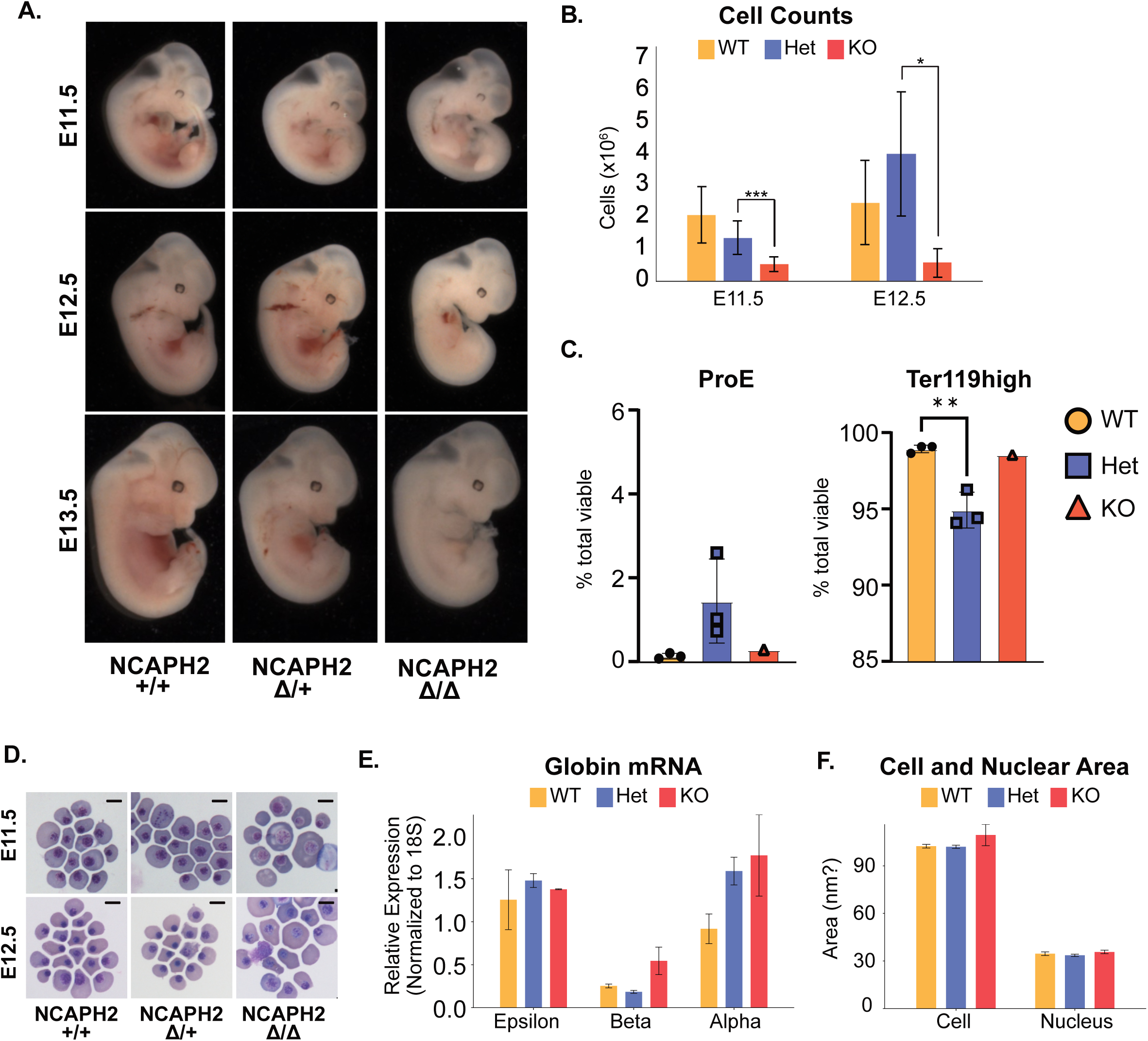
Ncaph2 Loss Disrupts Erythroid Maturation in vivo. (A) Pictures of *Ncaph2* 11/11 and control embryos at E11.5, E12.5 and E13.5. (B) Total peripheral blood live embryonic cell counts (E11.5 WT n=17, Het n=13, KO n=12, E12.5 WT n=5, Het n=3, KO n=4) (statistics determined from t test, * p-value <0.05, **p-value <0.01) (C) Flow cytometric analyses of indicated genotypes. Proerythroblasts (Pro-E) defined as CD71 positive, Ter119 negative. Example gating shown in Fig S1. (D) Cytospins of E11.5 and 12.5 primitive erythroblasts in *Ncaph2* 11/11, *Ncaph2* 11/+, and *Ncaph2* +/+ visualized using Wright-Giemsa staining. Scale bars = 10um (E) Quantitative analysis of globin expression in primitive erythroblasts using qPCR. (WT n=3, Het n=2, KO n=2) (F) Imaging flow cytometric analyses of cell and nuclear size.

Loss of Kmt5A, which deposits the Condensin II interacting mark H4K20me1, results in a block in erythroid differentiation associated with a dramatic loss of nuclear condensation (Malik et al., 2015). Surprisingly, erythroblasts from *Ncaph2* 11/11 embryos had nuclei similar in appearance to cells from littermate controls (Fig 1D). Further assessment of nuclear size by imaging flow cytometry showed comparable average nuclear sizes among the genotypes (Fig 1F), suggesting that unlike Kmt5a, Ncaph2 is dispensable for chromatin condensation during the terminal maturation of primitive erythroid cells, despite the essential role it plays in mitotic condensation.

### Ncaph2 and Kmt5aA Regulate a Shared set of Genes

To determine the effects of Ncaph2 loss on gene expression, we performed RNA-seq on erythroblasts from *Ncaph2* 11/11 and littermate control embryos (*Ncaph2* 11/+, and +/+) at E11.5. This analysis reveals that *Ncaph2* deletion results in 773 upregulated and 957 downregulated genes in erythroblasts (Fig 2A,B). The upregulated genes were enriched for terms related to translation, p53 pathway, and innate immunity (Fig 2C). The downregulated genes were enriched for multiple terms related to mitotic cell cycle and mitotic chromosome segregation (Fig 2C). In addition, all other members of the canonical Condensin II complex and the related Condensin I complex were significantly downregulated following Ncaph2 loss. (Fig 2D).

**Figure 2.**
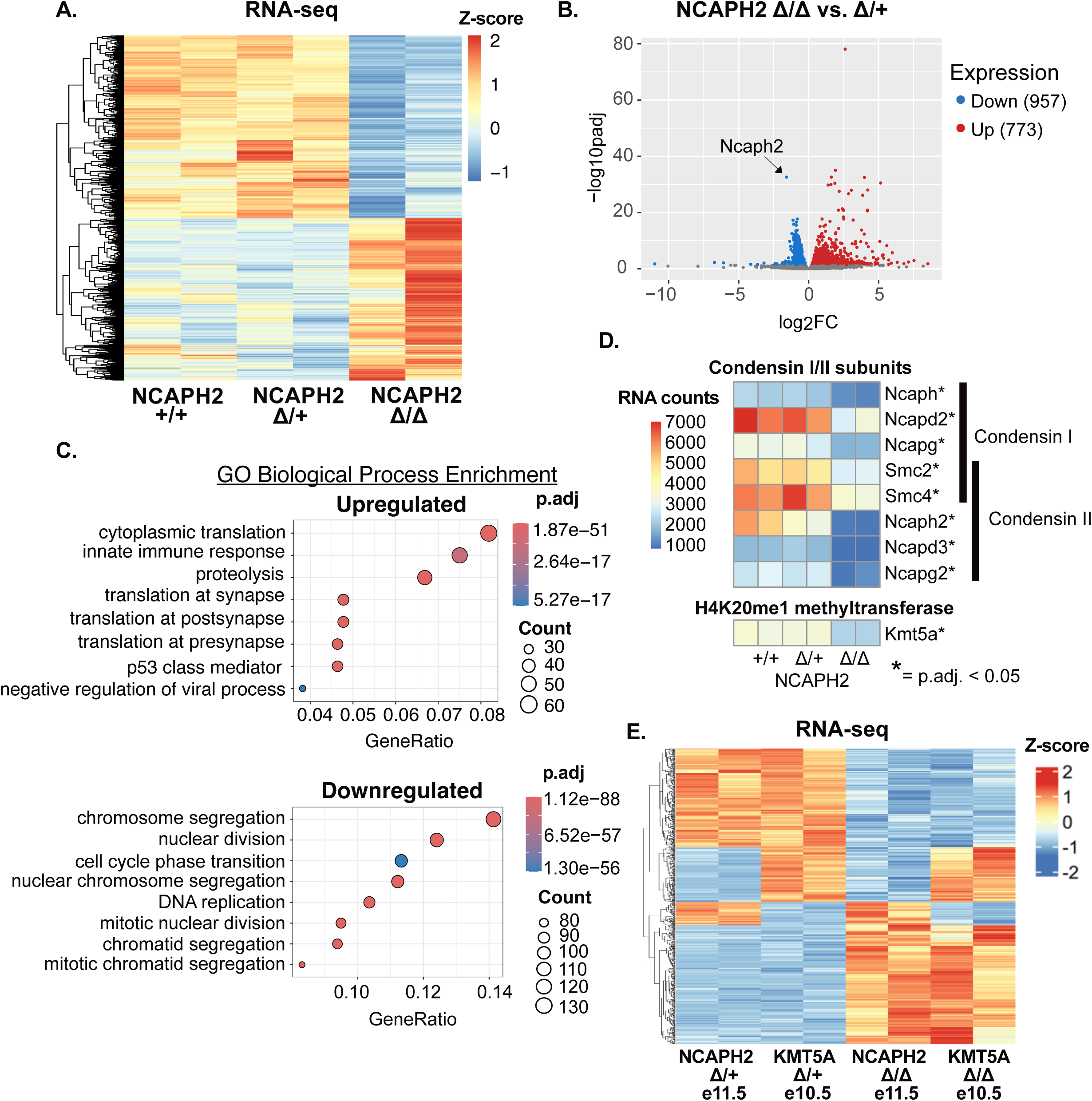
Loss of Ncaph2 alters erythroid gene expression. (A) Heat map of z-score normalized expression of differentially expressed genes (*Ncaph2* 11/11 vs. 11/+) for *Ncaph2* 11/11, 11/+, and +/+ erythroblasts at E11.5. N=2 for each genotype. (B) Volcano plot showing differential gene expression in *Ncaph2* 11/11 vs. 11/+ (C) Gene Ontology Biological Process Enrichment analyses of differentially upregulated and downregulated genes. (D) Heat map plotting z-score normalized expression of Condensin I and II complex subunits as well as Kmt5A in indicated genotypes. (E) Heatmap of genes differentially expressed in Ncaph2 (E11.5) and Kmt5a (E.10.5) mutant embryos. N=2 for each indicated genotype.

Ncaph2 is capable of recognizing H4K20me1 (Liu et al., 2010). Notably, upon loss of Ncaph2, we observed a significant decrease in levels of Kmt5a, the enzyme solely responsible for deposition of H4K20me1 (Fig. 2D). To investigate the relationship between regulation of gene expression downstream of Ncaph2 and Kmt5a, we compared the Ncaph2 RNA-seq analysis to a previous analysis we performed using mice harboring a conditional (EpoRCre) deletion of Kmt5A. (Malik et al., 2015) We found significant overlap among both up- and down-regulated genes in the two backgrounds (Figs 2E, S2). The shared set of upregulated genes was enriched for pathways related to immunity and inflammation (Fig S2), although this group also included several transcription factors that are typically silenced while erythroid cells mature, including Gata2, Hhex, and Jun. Pathway analysis of shared downregulated genes was significant for terms related to mitosis, chromatin segregation, and cell cycle (Fig S2). In contrast, comparison of the *Ncaph2* 11/+ to wild type littermate controls revealed few differences, suggesting a minimal effect of EpoRCre expression or heterozygous loss of *Ncaph2* (Fig S2). Together, these data suggest that Ncaph2 loss may affect some genes indirectly through changes in Kmt5a levels.

### Ncaph2 is Enriched at Promoters of Highly Expressed Genes Important for Erythroid Maturation and Cell Cycle Progression

To delineate the genes directly regulated by Ncaph2, we performed CUT&RUN (Skene and Henikoff, 2017) for Ncaph2 in wild type erythroblasts from E11.5 embryos. The majority of Ncaph2 occupancy (>67%) occurred at promoters, with a smaller number of Ncaph2 peaks in introns and distal intergenic regions (Fig 3A). Motif analysis established that regions of Ncaph2 occupancy were enriched for general transcription factors, such as SP1, as well several erythroid transcription factors, including GATA1:TAL1 and BACH1 (Fig 3B), while gene ontology analyses revealed enrichment for cell cycle related terms (Fig 3C). Intriguingly, levels of Ncaph2 occupancy were highly correlated with gene expression, with the most highly expressed genes having the highest levels of Ncaph2 occupancy (Fig 3D).

**Figure 3.**
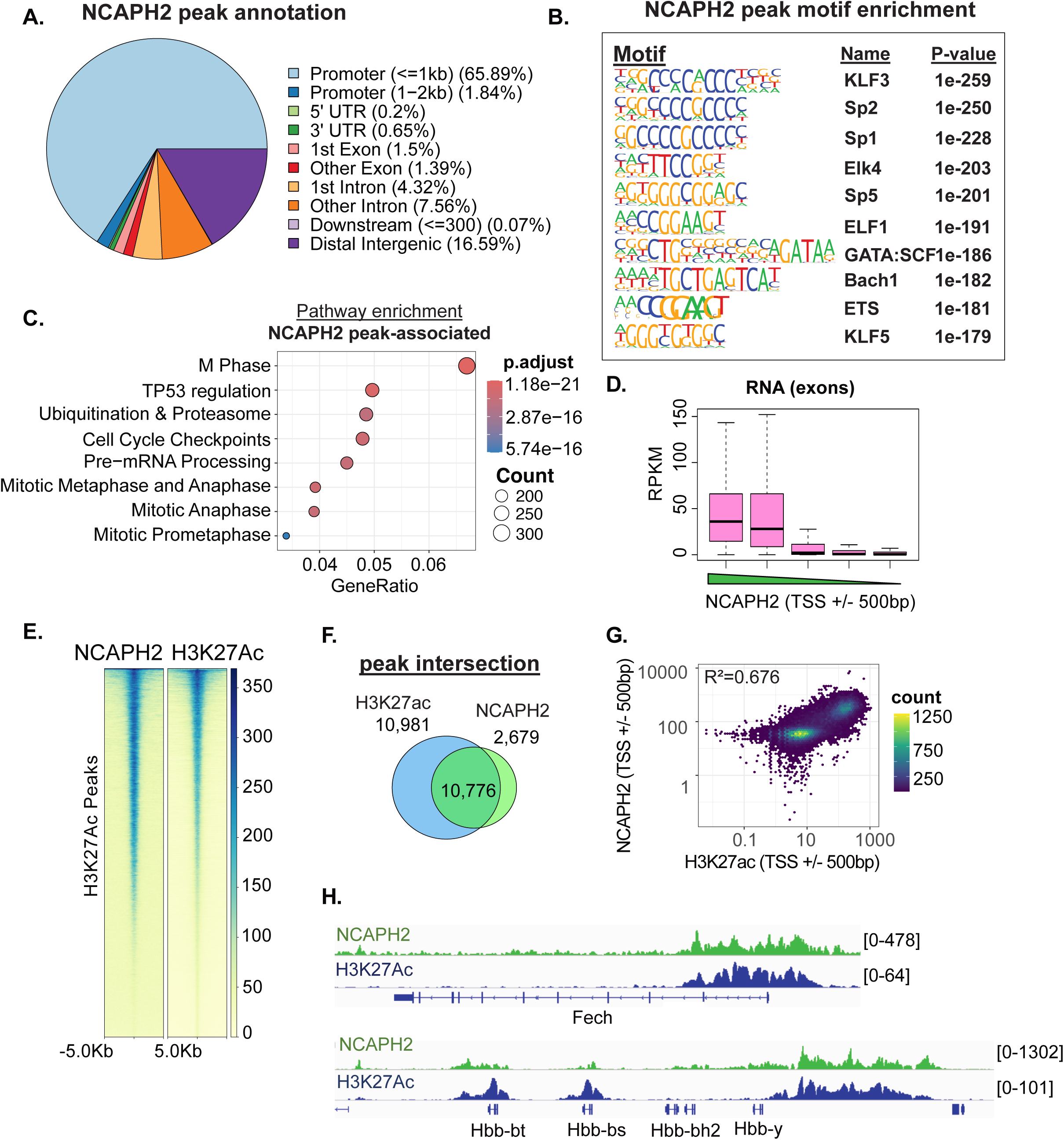
Ncaph2 occupies the promoters of highly expressed genes. (A) Genomic annotation of Ncaph2 peaks. (B) Motif enrichment analyses of Ncaph2 peaks (C) Pathway enrichment of genes associated with Ncaph2 occupancy (within 2kb of TSS). (D) Box plot showing average RNA RPKM over exons for all genes divided into quintiles based on Ncaph2 levels at promoters (TSS+/-500bp). (E) Heatmap showing the Ncaph2 enrichment over H3K27ac peaks in E14.5 fetal liver erythroid cells (F) Venn diagram showing intersection of H3K27Ac and Ncaph2 peaks. (G) Correlation scatter plot of H3K27Ac and Ncaph2 RPKM over promoters (TSS +/- 500bp). R2 represents Pearson correlation coefficient (H) Ncaph2 and H3K27ac occupancy at the Fech and beta globin loci. Ncaph2 data represent merged replicate RPKM (n=3). H3K27Ac data represent merged replicate RPKM from GSE132130 (n=2)

We next interrogated the chromatin landscape at regions of Ncaph2 occupancy. As Ncaph2 was associated with highly expressed genes, we first analyzed the association of Ncaph2 with acetylated histone H3 K27 (H3K27Ac), a histone mark associated with active promoters and enhancers. In erythroid cells, H3K27Ac was present primarily at promoter regions and at distal intergenic regions, which likely represent enhancers (Fig S3). We found that Ncaph2 occupancy was highly correlated with H3K27Ac at peaks and promoters (Fig 3E-G). Regions of Ncaph2-H3K27Ac co-occupancy were enriched for pathways related to cell cycle and chromosome organization (Fig S3). Many highly expressed erythroid genes were Ncaph2 targets, including Ankyrin1 (Ank1), Alpha Spectrin (Spta1), and Ferrochelatase (Fech) (Fig 3E, S3). Ncaph2 was also present at several well-described erythroid enhancers, including those associated with alpha and beta globin (Fig 3H, Fig S3) Together, these data demonstrate that Ncaph2 occupies the transcription start site and enhancers of highly expressed genes with key roles in cell erythroid maturation.

### H4K20me1 is Enriched in the Body of Ncaph2 Target Genes

As Ncaph2 and Kmt5A appeared to regulate a shared set of genes, we next compared H4K20me1 and Ncaph2 occupancy in erythroblasts from wild type E11.5 embryos. While Ncaph2 was present primarily at transcription start sites, H4K20me1 was present primarily over gene bodies (Fig 4A-C). Although they did not directly overlap, Ncaph2 and H4K20me1 occupancy were significantly correlated (Fig 4D), with many genes having Ncaph2 enrichment at the transcription start site and H4K20me1 enrichment over the gene body (Fig 4 D-E). Similar to Ncaph2, regions of H4K20me1 occupancy were enriched for pathways significant for terms related to cell cycle (Fig 4F). H4K20me1 occupancy also correlated with gene expression, with the most highly expressed genes having the highest level of H4K20me1 (Fig 4G). Multiple highly expressed erythroid genes had regions of Ncaph2 and H3K27Ac co-localization at the transcription start site, with enrichment for H4K20me1 over the gene body (Fig 4H, S4).

**Figure 4.**
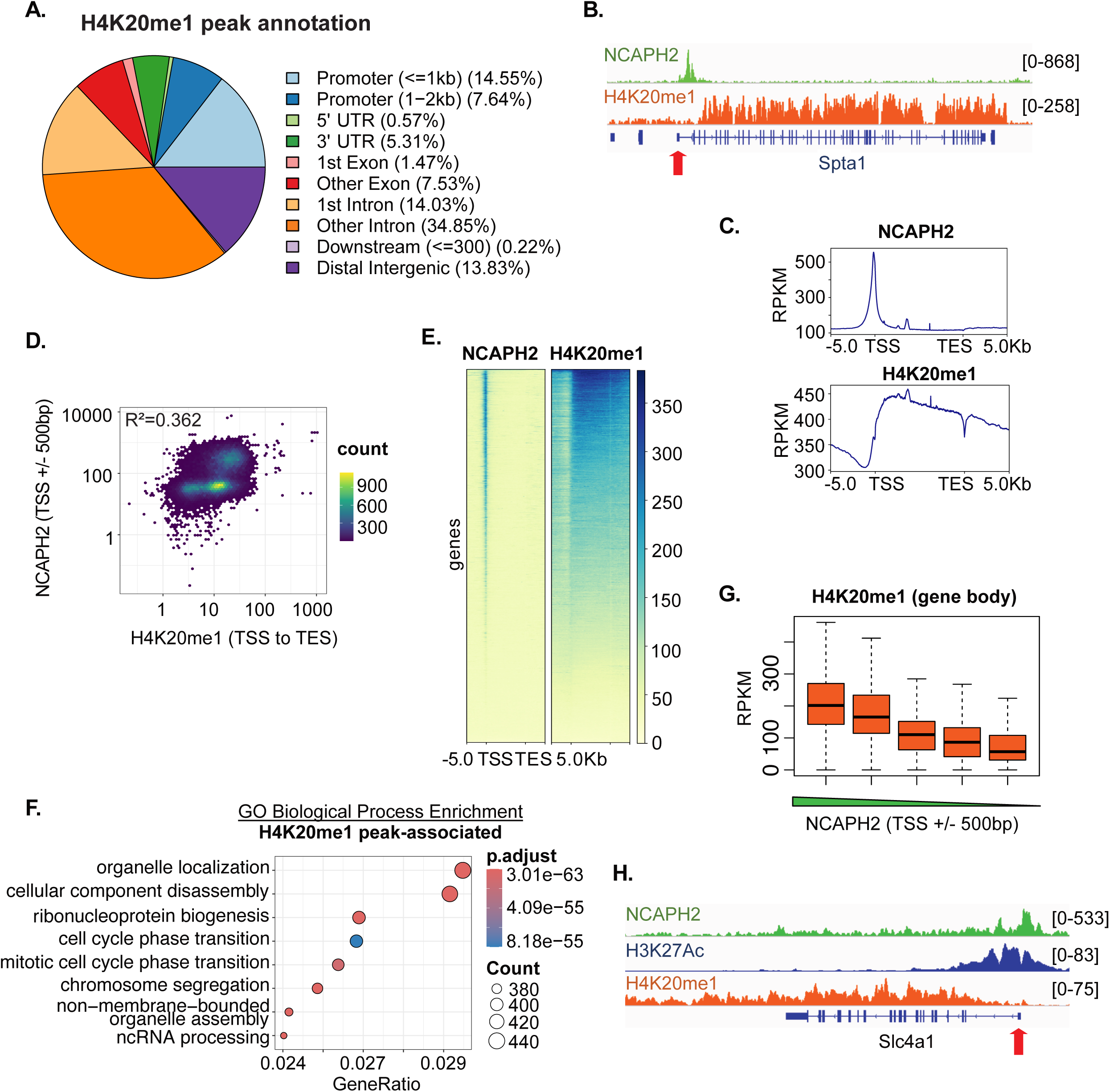
H4K20me1 is Enriched in the Body of Ncaph2 Target Genes. (A) Genomic annotation of H4K20me1 peaks. Data represent merged replicate RPKM (n=4). (B) Ncaph2 and H4K20me1 tracks at the Spta1 locus. Red arrow denotes promoter. Ncaph2 and H4K20me1data represent merged replicate RPKM (n=3 and 4, respectively). (C) Profile plots of Ncaph2 and H4K20me1 enrichment over genes (TSS to TES +/- 5kb) (D) Correlation scatter plot of H4K20me1 and Ncaph2 RPKM over gene-matched gene bodies (TSS to TES) and promoters (TSS +/- 500bp), respectively. R2 represents Pearson correlation coefficient (E) Heatmap showing the Ncaph2 and H4K20me1enrichment over genes (TSS to TES +/- 5kb) (F) Gene Ontology Biological Process Enrichment of genes associated with H4K20me1 peaks (G) Box plot showing average H4K20md1 RPKM over gene bodies (TSS to TES) for all genes divided into quintiles based on Ncaph2 levels at promoters (TSS+/-500bp). (H) Ncaph2 and H4K20me1 occupancy at the Slc4a1 loci. Red arrow denotes promoter. Ncaph2 and H4K20me1data represent merged replicate RPKM (n=3 and 4, respectively).

### Ncaph2 Target Genes are Downregulated Following Ncaph2 loss

To further interrogate the role of Ncaph2 in gene regulation, we correlated Ncaph2 occupancy in WT E11.5 erythroblasts with changes in gene expression following *Ncaph2* deletion. Many downregulated genes were directly associated with Ncaph2, while the subset of upregulated genes that were Ncaph2 target genes was smaller (Fig 5A, S5). Downregulated genes had significantly more Ncaph2 occupancy near their transcription start sites than upregulated genes (Fig 5B-C), and genes with the highest levels of Ncaph2 occupancy were the most downregulated in *Ncaph2* 11/11 erythroid cells (Fig 5D). Downregulated genes that were direct Ncaph2 targets include key erythroid genes such as Ank1, Klf1, and Spta1 (Fig S3,4). Pathway analyses of those genes was significant for terms related to cell cycle (Fig 5E).

**Figure 5.**
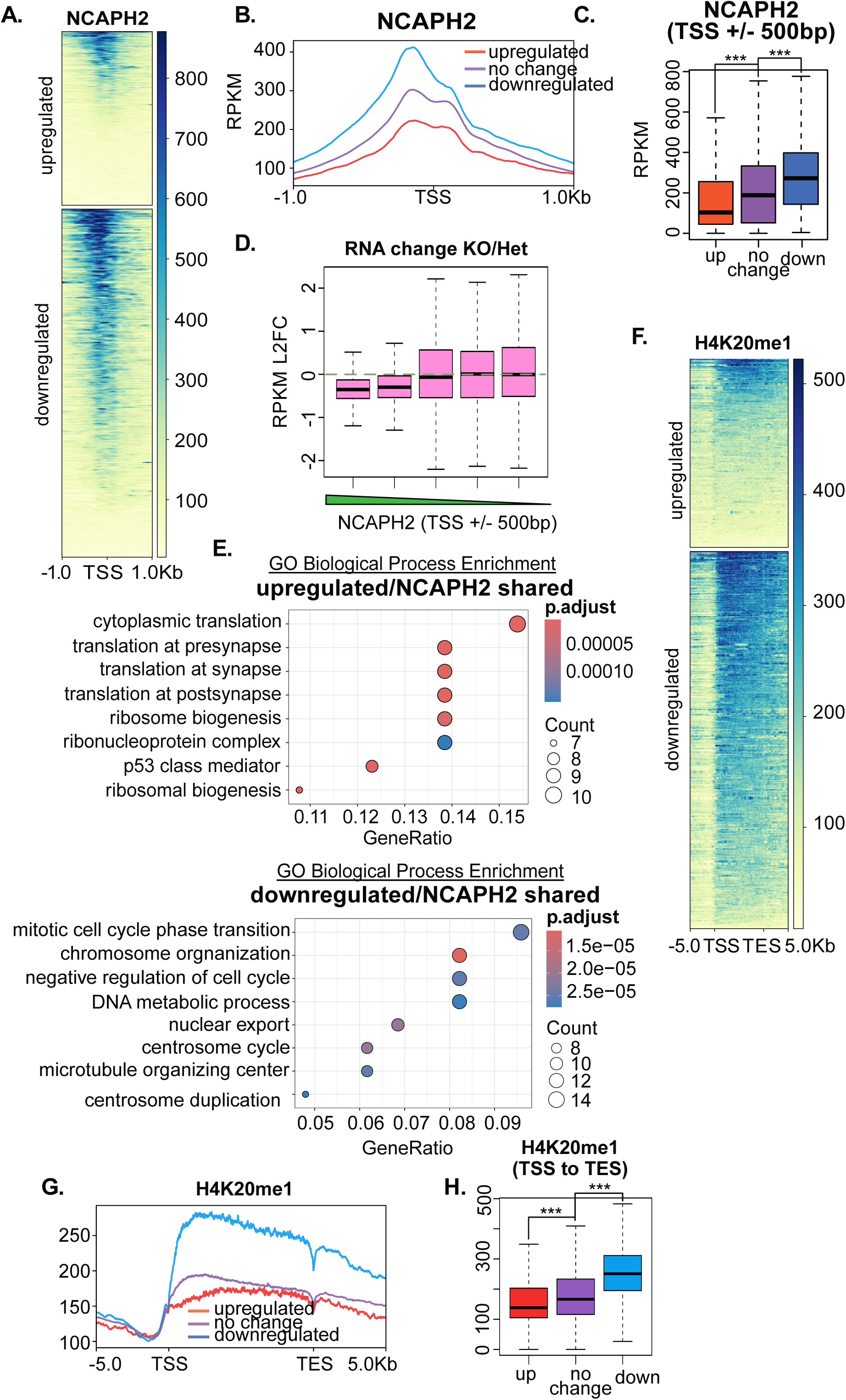
Ncaph2 promotes the expression of highly expressed erythroid genes. (A) Heatmaps of Ncaph2 and H4K20me1 over promoters (TSS +/- 1kb) of differentially expressed genes following Ncaph2 deletion. (B) Profile plot showing Ncaph2 levels at promoters of differentially upregulated, differentially downregulated, and not significantly changed genes following Ncaph2 deletion. (C) Box plot of Ncaph2 levels at the transcription start site of genes that are upregulated, downregulated, and do not change following Ncaph2 deletion. (statistics determined from t test, *** p-value < 2.2e -16) (D) Box plot showing L2FC RNA over exons for all genes divided into quintiles based on Ncaph2 levels at promoters (TSS+/-500bp). (E) Gene Ontology Biological Process Enrichment of differentially expressed genes that are direct Ncaph2 targets (peak within 2kb of TSS) (F) Heat map of H4K20me1 enrichment over genes differentially expressed upon Ncaph2 deletion. (G) Profile plot of H4K20me1 levels over genes (TSS to TES +/- 5kb) that are differentially upregulated, differentially downregulated, and not significantly changed following Ncaph2 deletion (H) Box plot demonstrating H4K20me1 levels at genes that are upregulated, downregulated, and do not change following Ncaph2 deletion.

In contrast, the subset of genes that were upregulated following *Ncaph2* deletion was enriched for terms related to translation and ribosome biogenesis (Fig 5E) and contained multiple ribosomal protein genes (Fig S5), which have been shown to be repressed by the Condensin II complex in other systems (Lancaster et al., 2021). In general, however, genes that increased expression upon *Ncaph2* deletion exhibited lower levels of Ncaph2 occupancy at their transcription start sites than genes with no change or decreased expression (Fig 5B), suggesting an indirect mechanism for many or most genes upregulated upon *Ncaph2* deletion.

Differentially expressed genes were also enriched for H4K20me1. The majority of downregulated genes had robust enrichment for H4K20me1 over the gene body (Fig 5F). In addition, downregulated genes had significantly higher levels of both Ncaph2 and H4K20me1 enrichment than upregulated genes (Fig 5G, H). Together these data suggest that in erythroid cells, Ncaph2 occupancy is primarily associated with promoting the expression of target genes.

### Ncaph2 Loss Increases H4K20me1 Levels in Maturing Erythroblasts

Our RNA-seq and CUT&RUN datasets, together with prior studies (Liu et al., 2010), suggest that Ncaph2 and H4K20me1 may work cooperatively to regulate gene expression. To further interrogate the relationship between the Condensin II complex and H4K20me1 in maturing erythroid cells, we assessed global levels of H4K20me1 in Ncaph2 mutant cells and littermate controls using immunofluorescence. Despite a decrease in the expression of Kmt5a mRNA, we identified a dramatic increase in H4K20me1 in the Ncaph2 mutant cells (Fig 6A,B). We therefore interrogated the expression of the other regulators of H4K20 methylation levels, including relevant methylases (Kmt5a, Kmt5b, Kmt5c) and demethylases (Phf8, Rad23a, Rad23b) and Hcfc1, a factor that can modulate H4K20me levels (Jorgensen et al., 2013). We did not observe large changes in the mRNA expression of any of these factors following *Ncaph2* deletion (Fig 6S), which is consistent with a large body of data suggesting Kmt5a levels are regulated primarily through posttranslational modifications, such as ubiquitination, in a cell cycle-dependent manner (Abbas et al., 2010, Centore et al., 2010, Oda et al., 2010). To further interrogate the relationship between Ncaph2 and Kmt5a levels, we performed immunofluorescence for Ncaph2 in E10.5 erythroblasts from *Kmt5a* fl/fl;EpoR Cre (*Kmt5a* 11/11) embryos. We observed a mild increase in Ncaph2 staining (Fig 6S), demonstrating an unexpected inverse relationship between Ncaph2 and H4K20me1 levels.

**Figure 6.**
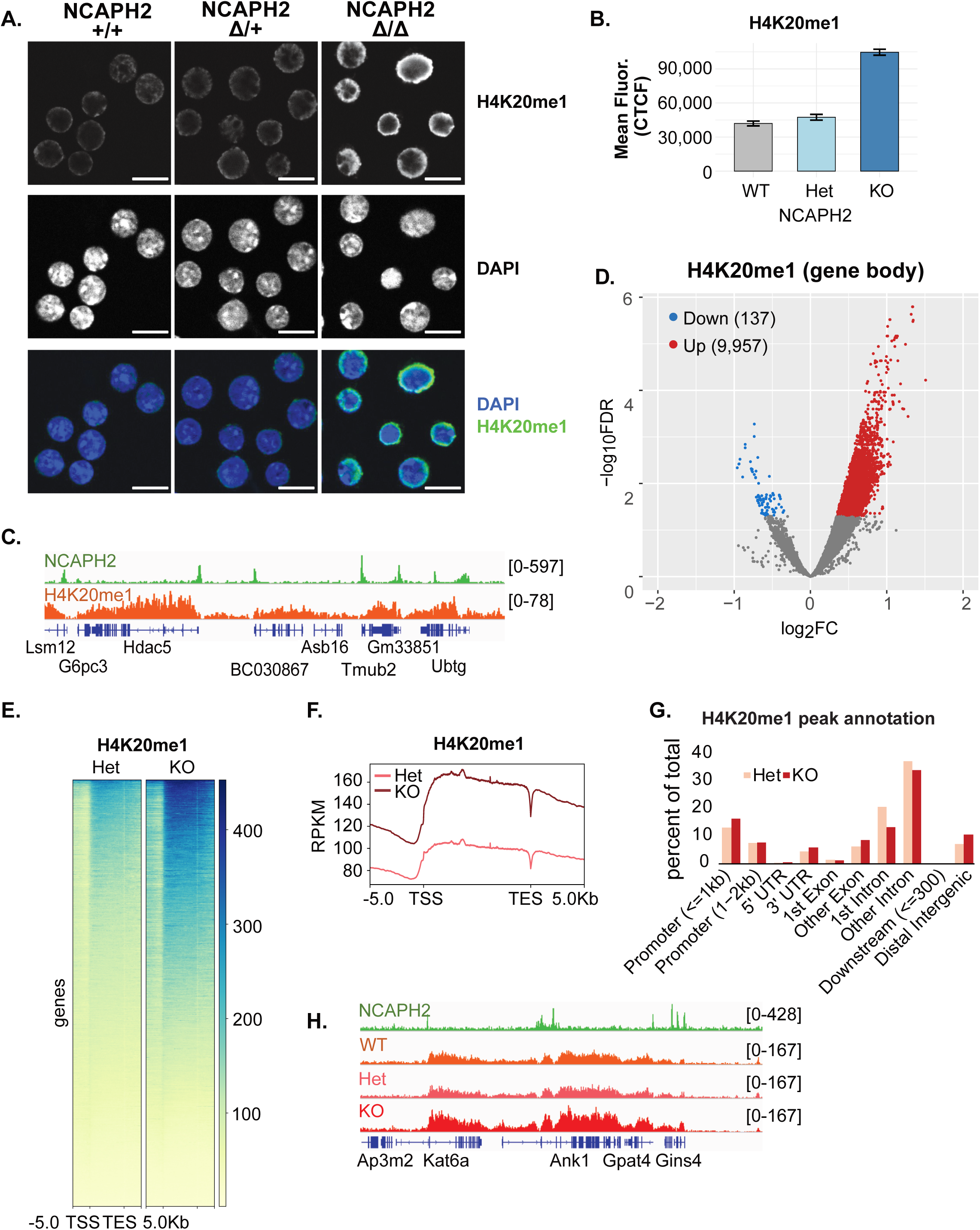
Ncaph2 Loss Increases H4K20me1 Levels in Maturing Erythroblasts. (A) Immunofluorescence staining for H4K20me1 and Ki67 in E11.5 erythroblasts of indicated *Ncaph2* genotype. (B) Quantification of H4K20me1 mean Correlated Total Cell Fluorescence (CTCF), quantification done in n=50 cells per Ncaph2 genotype, biological replicates (individual embryos) WT n=3, Het n=1, KO n=3 (C) Ncaph2 and H4K20me1 occupancy at locus containing several genes (D) Volcano plot comparing H4K20me1 in E11.5 erythroblasts from *Ncaph2* 11/11 and *Ncaph2* 11/+ embryos. (E) Heat maps showing H4K20me1 over genes (TSS to TES +/- 5kb) in erythroblasts from *Ncaph2* 11/11 and *Ncaph2* 11/+ embryos (F) Profile plot of H4K20me1 levels in El1.5 *Ncaph2* 11/11 and *Ncaph2* 11/+ (G) Genome annotations of H4K20me1 peaks in erythroblasts from *Ncaph2* 11/11 and *Ncaph2* 11/+ embryos (H) Ncaph2 and H4K20me1 occupancy at genes downregulated in *Ncaph2* 11/11 compared to *Ncaph2* 11/+.

Ncaph2 has a well-described role in the regulation of higher order chromatin structure, (Hoencamp et al., 2021), and at several loci Ncaph2 appeared to mark the boundary of regions of H4K20me1 occupancy (Fig 4C). We next sought to understand whether Ncaph2 loss affected the genomic distribution of H4K20me1 by performing CUT&RUN for H4K20me1 in *Ncaph2* 11/11 cells. Consistent with the immunofluorescence results, we observed a global increase in H4K20me1 occupancy in the Ncaph2 mutant cells, with 2884 genes that gained H4K20me1 occupancy compared to 182 genes that lost it (Fig 6D-F). Although *Ncaph2* 11/11 cells had higher levels of H4K20me1 at individual binding sites, the distribution of H4K20me1 occupancy was similar between *Ncaph2* 11/11 and control cells and we did not observe “spreading” of regions of H4K20me1 occupancy or major changes in genomic annotation following *Ncaph2* deletion (Fig 6G,H S6). Together, these data suggest that loss of the Condensin II complex increases the magnitude of H4K20me1 occupancy, but that the Condensin II complex does not govern its distribution.

### Ncaph2 Regulates the Expression of Genes Essential for Erythroid Cell Cycle Progression

Ncaph2 has a well-characterized role in cell cycle progression, and our studies identified global increases in H4K20me1, a histone mark whose level is highest during the G2/M phase of the cell cycle, following *Ncaph2* deletion. In addition, further analyses of the RNA-seq data revealed that many genes important for cell cycle progression were downregulated in *Ncaph2* 11/11 erythroblasts (Fig 7A). Consistent with these findings, BRDU incorporation revealed that the Ncaph2 mutant cells had abnormalities in cell cycle progression, with a build-up of cells in the G2/M phase of the cell cycle (Fig 7B,), and *ex vivo* culture of mutant erythroblasts revealed a defect in proliferation (Fig S7).

**Figure 7.**
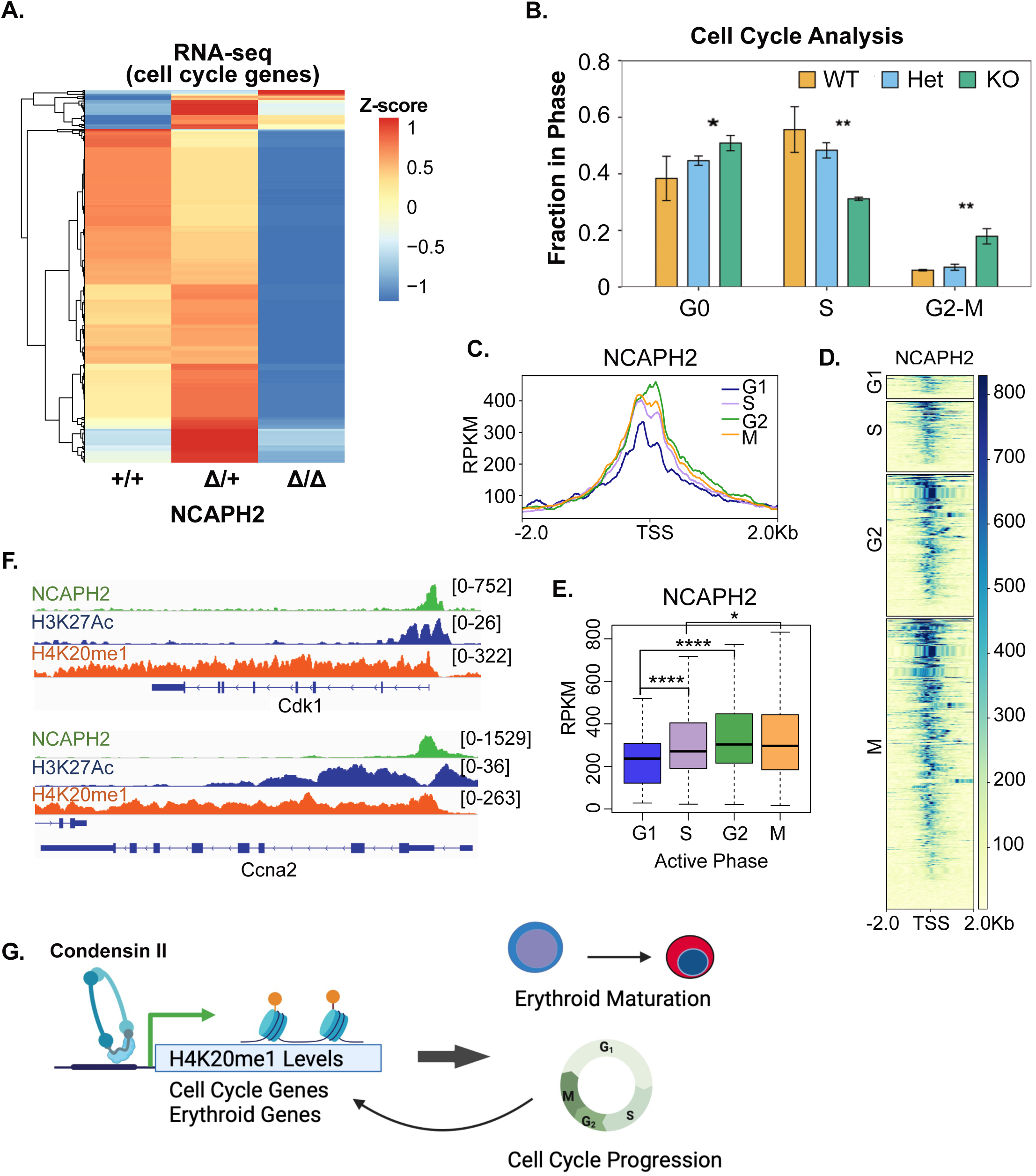
Ncaph2 Regulates the Expression of Genes Essential for Erythroid Cell Cycle Progression. (A) Differentially expressed cell cycle genes in *Ncaph2* 11/11 and *Ncaph2* 11/+ erythroblasts. (B) Cell cycle analyses of E11.5 erythroblasts of indicated genotype performed using BRDU staining. Statistics determined from t test, ** p-value < 0.01, * p-value < 0.05 (C) Profile plot of Ncaph2 occupancy at genes regulating indicated phases of the cell cycle in wild type E11.5 erythroid cells. (D) Heatmap of Ncaph2 over promoters (TSS +/- 2kb) of genes regulating indicated phases of the cell cycle in wild type E11.5 erythroid cells (E) Levels of Ncaph2 occupancy at genes involved in specific stages in of the cell cycle in wild type E11.5 erythroid cells. (F) Ncaph2 and H4K20me1 occupancy at Cdk1, Ccna2, and Ccnb2, key genes for progression through G2/M stage of the cell cycle. (G) Model of Ncaph2 and H4K20me1 activity during erythropoiesis.

In wild type erythroblasts, sites of Ncaph2 occupancy were significantly enriched for multiple pathways related to cell cycle progression. We therefore interrogated the hypothesis that Ncaph2 directly regulates genes that promote erythroid cell cycle progression. We identified Ncaph2 occupancy at the transcription start site of multiple genes that regulate cell cycle progression, with the highest level of occupancy at genes that regulate G2/M transition (Fig 7C-D) Many of these genes were significantly downregulated following *Ncaph2* deletion, including the key G2/M regulators Cdk1, ccna2 and ccnb2 (Fig 7F, S7). Together these data suggest that in addition to its well-established role in promoting mitotic chromatin condensation, the Condensin II complex may have a direct role in regulating the expression of genes that promote progression through G2/M.

## Discussion

Efficient erythropoiesis requires dramatic changes in gene expression in the setting of a rapidly dividing cell with a nucleus that is undergoing profound condensation prior to enucleation. Our data demonstrate an essential role for the Condensin II complex subunit Ncaph2 in erythropoiesis, with erythroid deletion of *Ncaph2* resulting in anemia and embryonic lethality. We show that the Condensin II complex is essential for erythroid cell cycle progression, with *Ncaph2* 11/11 erythroblasts accumulating in G2/M, similar to Condensin II disruption in other cell types (Macdonald et al., 2022). The variation in cell size observed in the *Ncaph2* 11/11 erythroblasts likely reflects this disrupted cell cycle progression. Previous studies have shown that the Condensin II complex promotes G2/M progression by binding modified histones and promoting mitotic chromatin condensation, with CDK1-mediated phosphorylation of Ncapd3 required for Condensin II activity (Abe et al., 2011). Our data further suggest that Ncaph2 directly regulates the expression of genes with critical roles in the G2/M phase of the cell cycle, including Cdk1, Ccnb2, and Ccna2. Notably the expression of these genes is decreased following *Ncaph2* deletion. Together our *in vivo* functional and genomic data suggests the Condensin II complex plays a more intricate role in cell cycle progression than previously realized.

Loss of the Condensin II complex results in dysregulated erythroid gene expression, and this effect appears to be at least partly a direct one. Ncaph2 is present at the promoters of genes that are highly expressed in erythroid cells, and gene expression levels correlate with Ncaph2 signal in the CUT&RUN assay. Moreover, deletion of *Ncaph2* results in decreased expression of these associated genes. Ncaph2 targets include genes essential for erythropoiesis, such as the membrane protein genes Ankyrin (Ank1), Alpha Spectrin (Spta1), Beta Spectrin (Sptb1), and Band 3 (Slc4a1), and the transcription factor Klf1, as well as the other components of the Condensin II complex, and the related complex Condensin I. Similar to studies in mESCs and HEK293 cells which identified Ncaph2 occupancy at promoters enriched for H3K4me3 and H3K27Ac(Yuen et al., 2017), we found extensive overlap between Ncaph2 and H3K27Ac occupancy. The Condensin II complex has also been found at super enhancers in murine embryonic stem cells, and is important for the expression of their target genes (Dowen et al., 2013). Ncaph2 was highly enriched at key erythroid enhancers, including the beta globin locus control region (LCR) and the alpha globin enhancers. Surprisingly, however, the expression of the globin genes was not affected by Ncaph2 loss. The study of Ncaph2 function at enhancers will be an intriguing area for further study.

The Ncapd3 and Ncapg2 subunits of the Condensin II complex can recognize H4K20me1, and the accumulation of H4K20me1 during early mitosis has been demonstrated to facilitate loading of the Condensin II complex during mitosis (Liu, Nature, 2011). Previous work from our group demonstrated a critical role for the histone methyltransferase Kmt5a and its histone mark H4K20me1 in erythroid chromatin condensation (Malik et al., 2015), but surprisingly the Condensin II complex appears to be dispensable for maturation-related chromatin condensation during primitive erythropoiesis, since *Ncaph2* 11/11 erythroblast nuclei are similar to wildtype controls in size and appearance. These data suggest that the mechanism of chromatin condensation that erythroid cells use during terminal maturation is distinct from that which occurs during mitosis.

The relationship between the Condensin II complex and the H4K20me1 histone mark in erythroid cells appears to be complex. In contrast to M-phase synchronized Hela cells(Liu et al., 2010), in erythroid cells H4K20me1 and Ncaph2 exhibit distinct patterns of genomic occupancy in maturing erythroblasts. While both Ncaph2 and H4K20me1 are present at highly expressed genes, Ncaph2 is found primarily at transcription start sites, while H4K20me1 occurs primarily over gene bodies. Several other studies have identified H4K20me1 over the bodies of actively transcribed genes, with the level of H4K20me1 occupancy correlating with the level of transcription. (Shoaib et al., 2021, Barski et al., 2007) The distributions we observe in erythroid cells suggest that loading at gene promoters of Ncaph2, and thus by extension the Condensin II complex, does not involve the H4K20me1 histone mark.

We observe an inverse relationship between Ncaph2 and H4K20me1 levels, with loss of Ncaph2 leading to a substantial increase in H4K20me1 levels. Somewhat paradoxically, Kmt5a expression is downregulated following *Ncaph2* deletion. While Kmt5a expression levels may decrease, however, they are not quantitatively eliminated, and given the stability of histone methylation, high levels of the H4K20me1 mark are still easily accomplished, especially given that expression levels of the demethylase that would otherwise erase this mark, PHF8, are relatively low in erythroid cells independent of genetic background. In addition, we observe accumulation of erythroid cells in G2/M phase upon *Ncaph2* deletion. H4K20me1 levels are normally cell cycle dependent, peaking in G2/M (Beck et al., 2012), and so the erythroid cell population as a whole would therefore be expected to exhibit higher levels of this mark. While it is possible that Ncaph2 directly regulates H4K20me1 levels, our current data favor a model where the reciprocal nature of their levels reflects their participation in distinct pathways, potentially linked through cell cycle

### Data Availability

Data generated in this study are available under Geo accession numbers GSE267278 (RNA-seq) and GSE267313 (CUT&RUN).

Other Data sets used in this study include: Setd8 KO RNA-seq e10.5 mouse erythroblasts: GSE83809 (Malik et al., 2015), and H3K27Ac ChIPseq e14.5 mouse fetal liver: GSE132130.(Fox et al., 2020)

**Figure S1.**
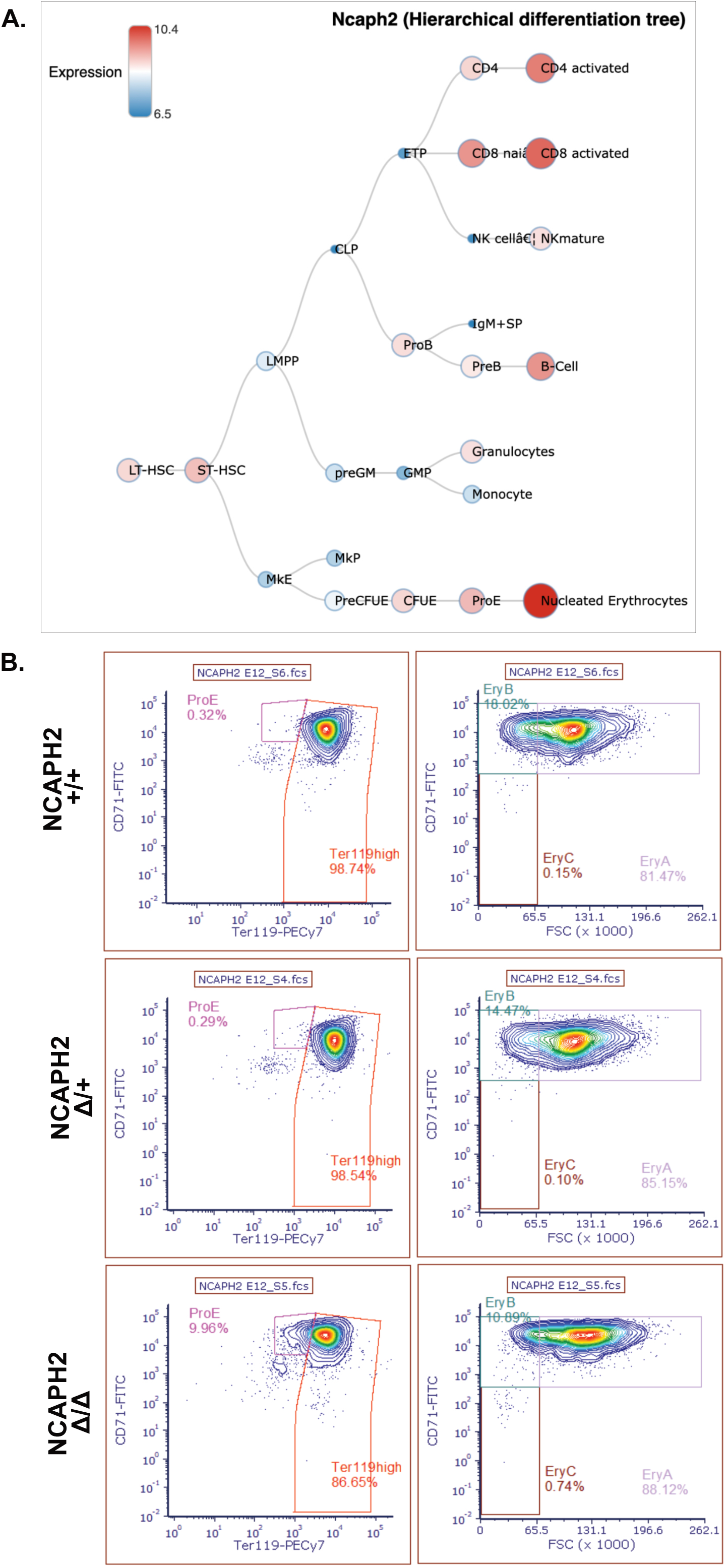
(A) Gene expression of Ncaph2 from blood spot data base (bloodspot.eu, (Bagger et al., 2019)) (B) Representative flow plot for each genotype showing CD71 vs Ter119 and CD71 vs FSC

**Figure S2.**
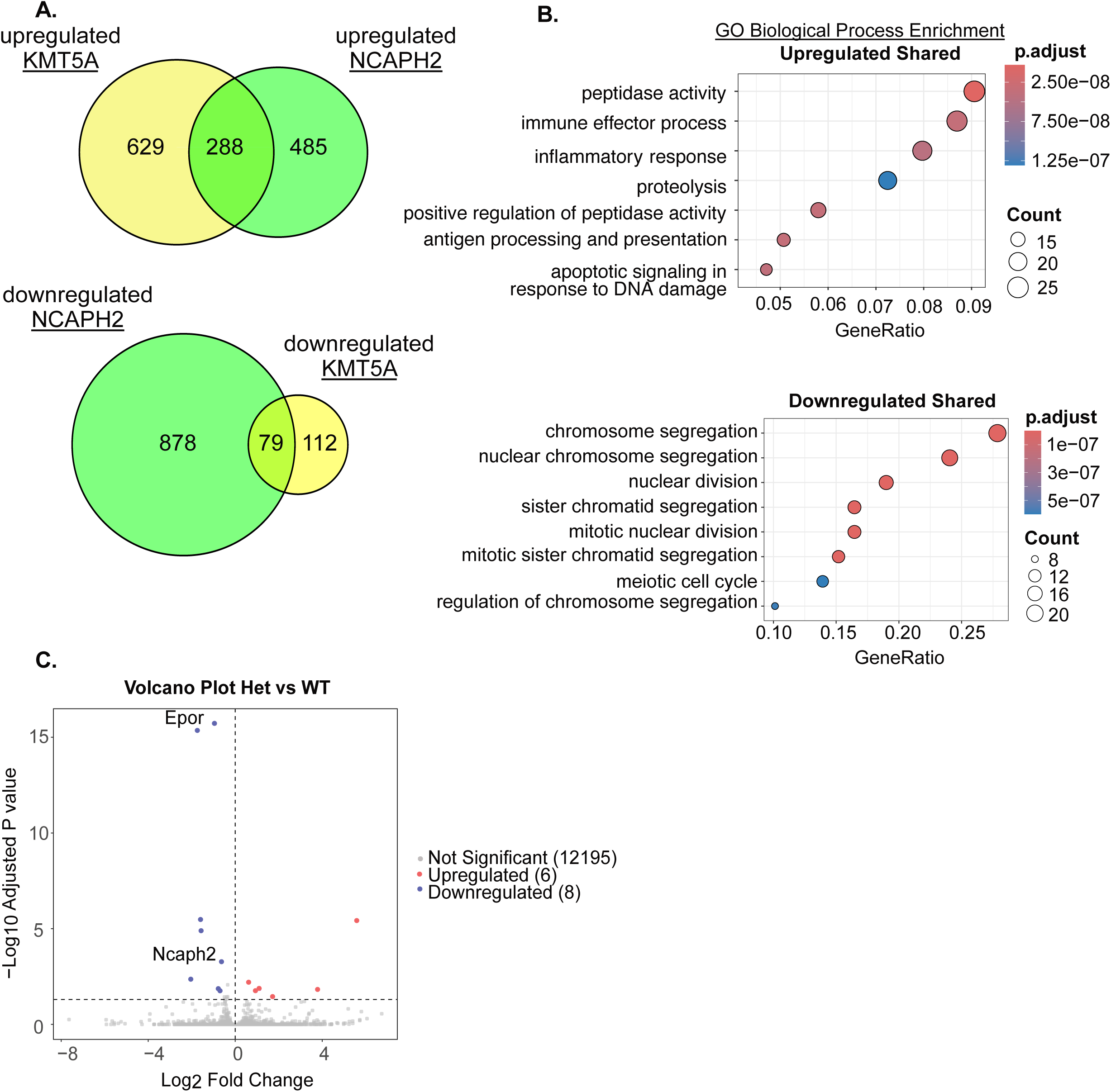
(A) Intersection of differentially expressed genes in *Ncaph2* 11/11 vs. Δ/+ and *Kmt5A* 11/11 vs. Δ/+ (B) Gene Ontology Biological Process Enrichment analyses of shared subsets (upregulated and downregulated) of differentially expressed genes (C) Volcano plot showing differential gene expression in *Ncaph2* 11/+ (EpoRCre +) vs. WT (EpoRCre -)

**Figure S3.**
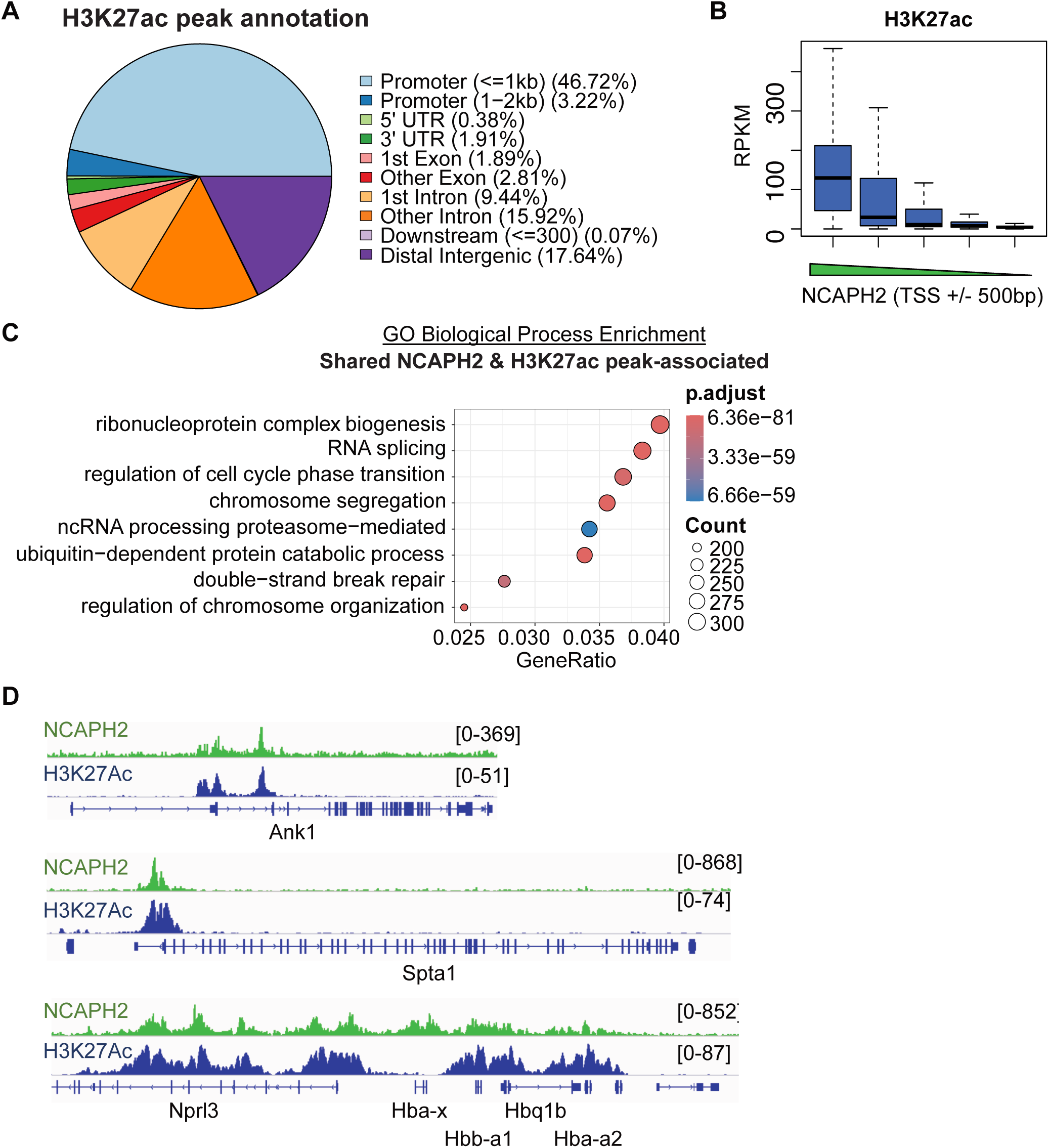
(A) Genomic annotation of H3K27Ac peaks (B) Box plot showing average H3K27Ac over promoters (TSS +/-500bp) for all genes divided into quintiles based on Ncaph2 levels at promoters (TSS+/- 500bp). (C) Gene Ontology Biological Process Enrichment analyses of genes associated with shared Ncaph2 and H3K27Ac occupancy (within 2kb of TSS). (D) Ncaph2 and H3K27Ac occupancy at Spta1, Ank1, and beta-globin loci. Ncaph2 data represent merged replicate RPKM (n=3). H3K27Ac data represent merged replicate RPKM from GSE132130 (n=2)

**Figure S4.**
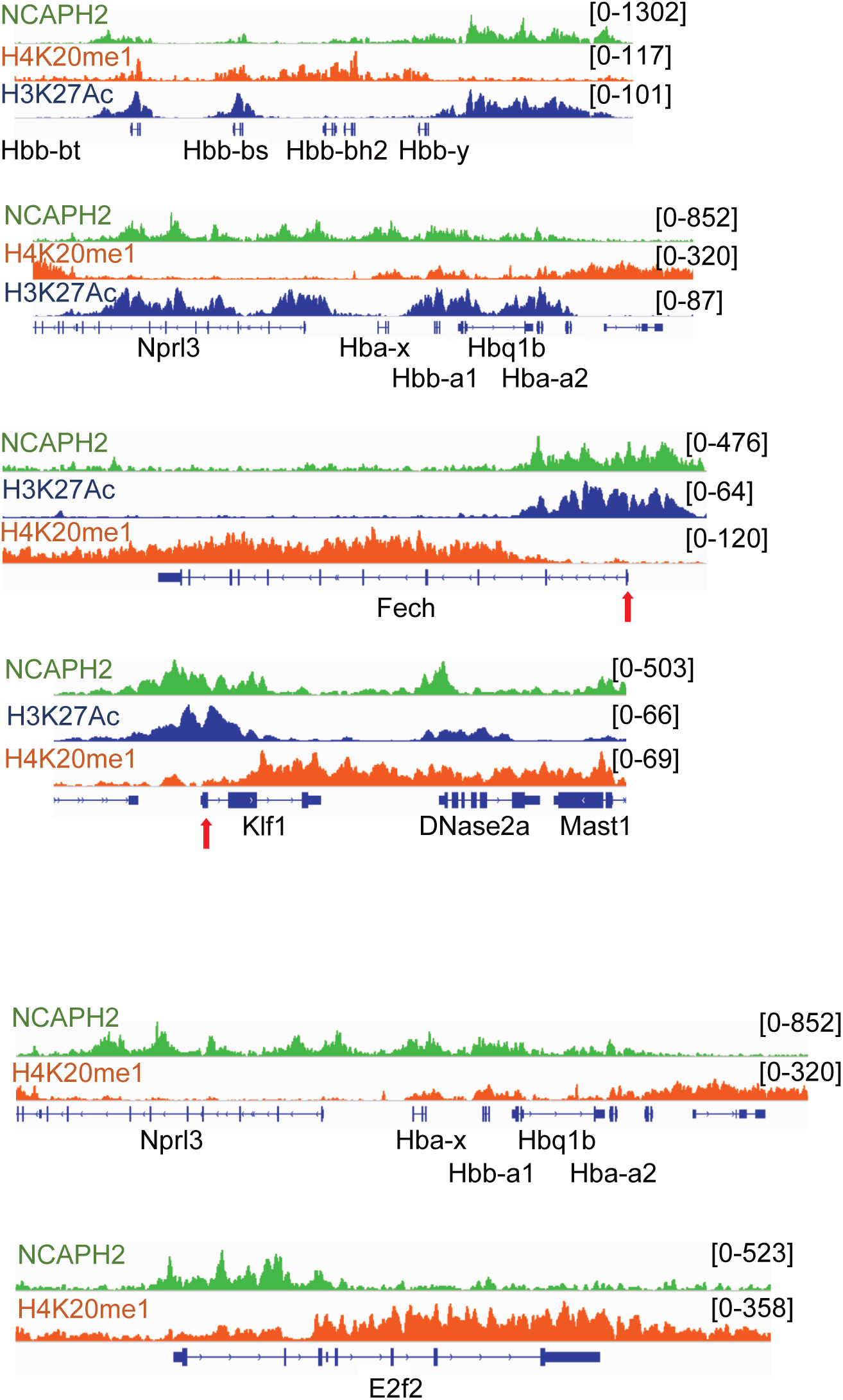
Ncaph2, H4K20me1, and H3K27ac occupancy at the beta-globin, alpha-globin, FECH, and Klf1 loci. Ncaph2 and H4K20me1data represent merged replicate RPKM. n=3 Ncaph2 n= 4 H4K20me1. N=2 H3K27Ac

**Figure S5.**
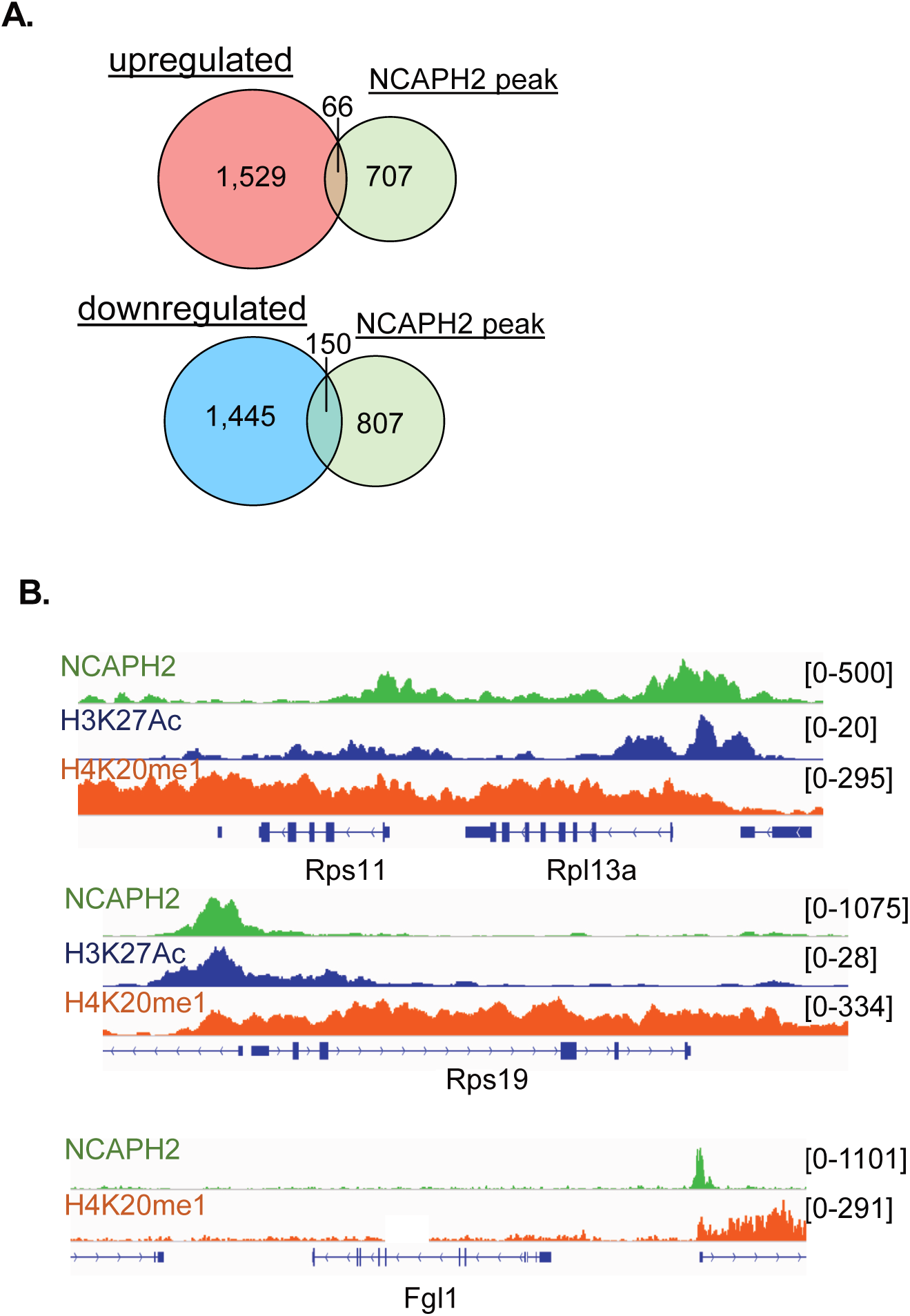
(A) Intersection of differentially expressed and Ncaph2 peak-associated (within 2 kb TSS) genes (B) Ncaph2, H4K20me1, and H3K27ac occupancy at various Rps and Fgl1 loci

**Figure S6.**
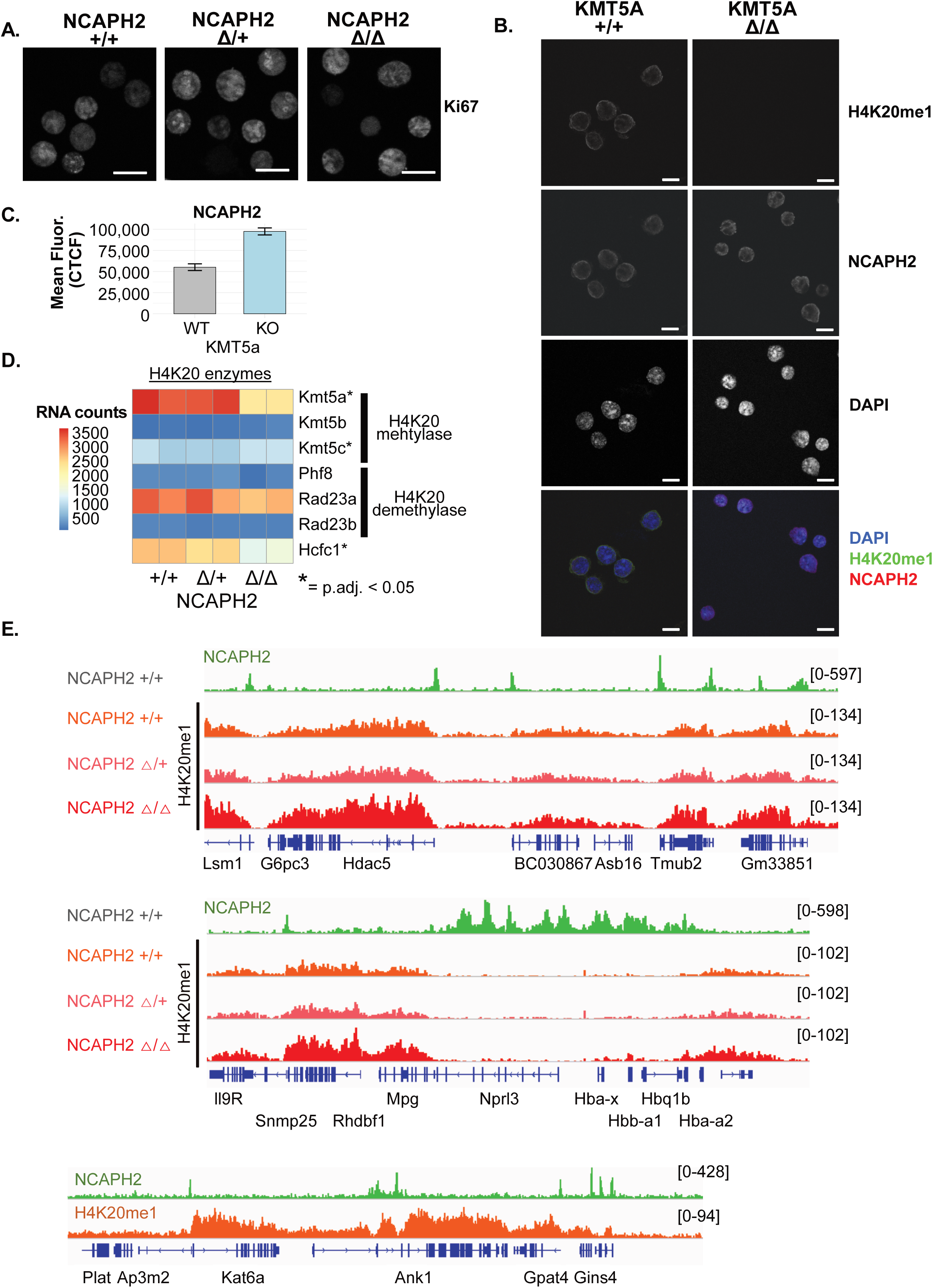
**(A)** Immunofluorescence for Ncaph2 in E10.5 erythroblasts of indicated *Kmt5a* genotype. **(B)** Quantification of Ncaph2 mean Correlated Total Cell Fluorescence (CTCF), n=50 cells per *Kmt5a* genotype **(C)** Ncaph2 and H4K20me1 occupancy at several loci for indicated NCAPH2 genotypes

**Fig S7.**
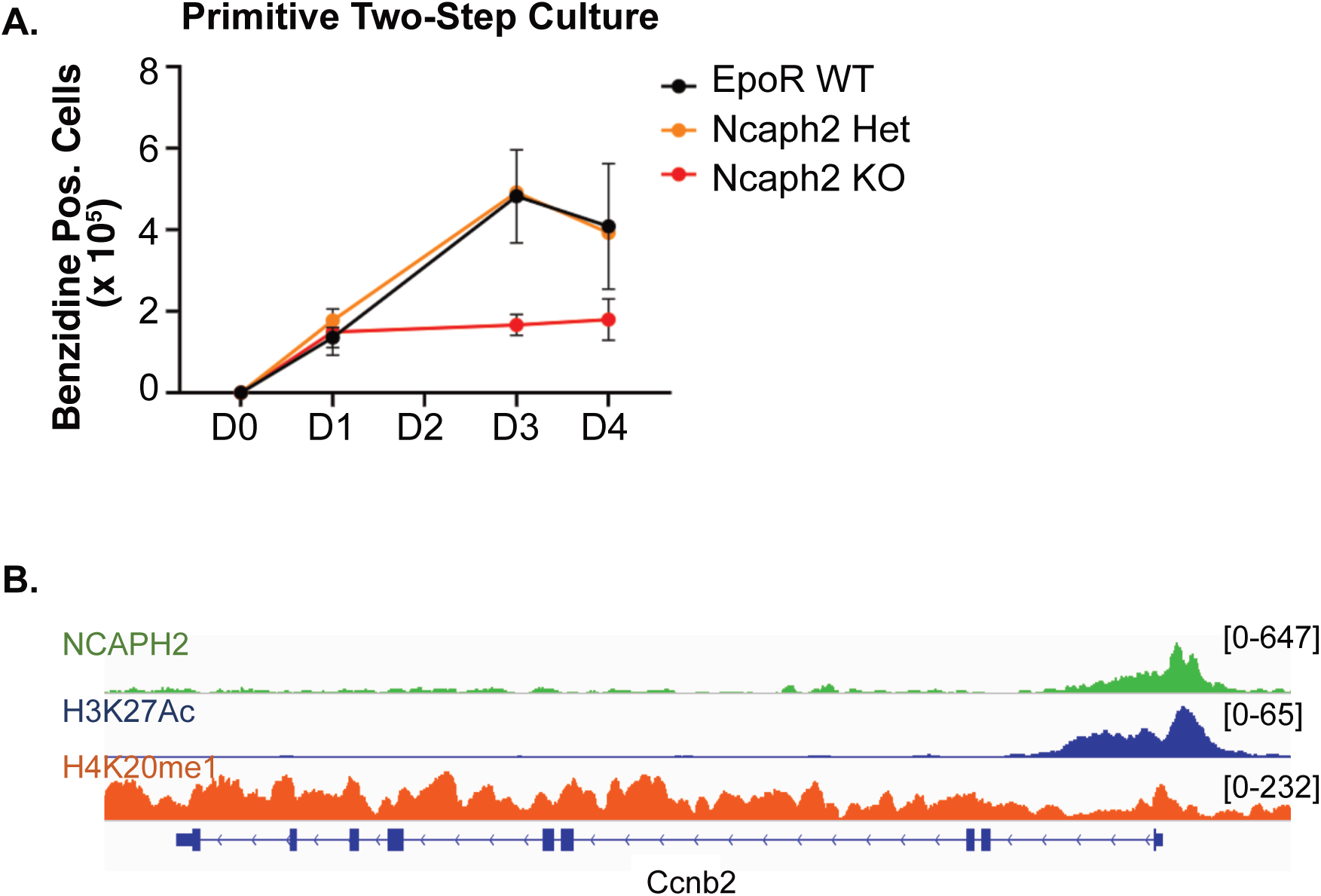
(A) Two-step culture of primitive erythroblasts of indicated genotypes. (B) Ncaph2, H4K20me1, and H3K27ac occupancy at Ccnb2 locus

## Methods

### Generation of Timed Embryos

Experiments involving mice were conducted following approval by the University of Rochester’s Committee on Animal Resources. *Ncaph2* Fl/Fl females were paired with *Ncaph2* Fl/+ Cre+/- males to produce *Ncaph2* 11/11, *Ncaph2* 11/+, and *Ncaph2* +/+ embryos. Vaginal plugs marked embryonic day 0.5 (E0.5), and pregnant dams were euthanized via followed by dissection of the embryos for further analysis. Peripheral blood was isolated as described previously (Kingsley et al., 2004).

### Cytospin Preparation and Analysis

Erythroblasts were isolated from peripheral blood and counted, with 100,000 cells prepared for cytospin analysis. Cells were pelleted, resuspended in SLAT-H (PB2, BSA, Heparin) media, and briefly incubated. The cytospin assembly was prepared, and the cell sample was applied and spun at 400xg for 3 minutes. Following centrifugation, the slides were air-dried and stained with Wright-Giemsa. For mounting, permount/xylene solution was applied, coverslips were placed, and the slides were air-dried overnight. Images were taken using a Nikon Eclipse 80i microscope equipped with a Nikon DS-Fi1 camera and analyzed using NIS-Elements software.

### Flow cytometry

For maturation analysis, peripheral blood samples were strained and washed with PB2 (PBS supplemented with CaCl_2_, Glucose, BSA). Samples were incubated for 10 minutes on ice with normal rat serum and incubated for 20 minutes on ice with a staining mix of PECyanine7-Ter119 (BD Pharmingen, Cat# 5557853) and FITC-CD71 (Invitrogen, Ref# 11-0711-82). Samples were washed again with PB2 and further stained with DAPI (Biolegend Cat# 422801) and DRAQ5 (Invitrogen Ref#65-0880-96) before analysis with the BD LSR Fortessa flow cytometer. Flow data acquisition was carried out using FACSDiva and further analyzed by FCS Express 7 – Research edition. The lasers used were RedC for DRAQ5, VioletF for DAPI, YellowGreenA for PE-Cyanine7, and BlueB for FITC, and FSC/SSC. The gating scheme involved exclusion of aggregates and cell debris with FSC/SSC, followed by gating viable cells using DAPI, and exclusion of nucleated maternal cells by DRAQ5. The viable cells were then gated based on Ter119vsCD71. The Ter119high population was further characterized as EryA, EryB, EryC based on FSC as described in (Koulnis et al., 2011).

For cell cycle analysis, cells were labeled with BrdU by resuspending them in IMDM medium containing Epo and BrdU, followed by incubation at 37°C. Surface staining was performed using PE Ter119, FITC CD71, and PECy7 CD117 antibodies on ice. Cells were washed, fixed with BD Cytofix/Cytoperm buffer, and subjected to intracellular staining with BD Perm Buffer Plus. DNase treatment was applied to allow for BrdU staining, followed by incubation with APC-BrdU antibody. The prepared cells were analyzed using ImageStreamX with filter settings of 405-50 for the blue channel (DAPI), 488-200 for the green channel, and 658-120 for the red channel. For analyses of cell and nuclear size, erythroblasts were stained with anti-CD71 (eBioscience), anti-Ter119 (eBioscience), and DRAQ5 (eBioscience) and run on the ImageStream X (Amnis).

### Immunofluorescence Staining and Analysis

Erythroblasts were isolated from embryos and prepared for cytospin analysis. Following cytospin preparation, cells were fixed in 4% paraformaldehyde for 20 minutes at room temperature and then washed three times with 1X PBS. Permeabilization was performed using 0.1% Triton X-100 in 1X PBS for 5 minutes on ice, followed by three washes with PBS. Blocking was conducted using 5% BSA in PBS-T for 1 hour at room temperature. Cells were then incubated with primary antibodies (Ncaph2: ab200659, H4K20me1: Invitrogen MA5-18067) diluted 1:200 in blocking buffer overnight at 4°C. After washing with PBS-T, secondary antibodies (Invitrogen Alexa Fluor 488: A-11001, Alexa Fluor 647: A-21235) diluted 1:500 in blocking buffer were applied for 1 hour at room temperature. Cells were washed with PBS-T and stained with DAPI diluted 1:1000 in PBS-T for 1 minute in the dark. Finally, slides were mounted with ProLong Gold antifade reagent with DAPI. Imaging was performed using a Nikon A1R Laser Scanning Confocal and TIRF microscope. To quantify cell intensity, the integrated density of each cell was measured using ImageJ’s “measurement” function. For each condition, 50 regions of interest (ROIs) were selected, with a background ROI drawn outside the cells for comparison. The corrected total cell fluorescence (CTCF) was then calculated using the formula: CTCF = integrated density in ROI − (area of ROI × mean intensity of background). This method corrects for background fluorescence, providing an accurate measure of the cell’s fluorescence intensity, and was based on the protocol “Measuring cell fluorescence using ImageJ” by J Hammond. This protocol is available online: https://theolb.readthedocs.io/en/latest/imaging/measuring-cell-fluorescence-using-imagej.html (Accessed October 2024).

### Genomics

RNA was extracted using the Qiagen RNeasy Kit. RNA library preparation and sequencing were performed by the University of Rochester Genomics Research Center. CUT&RUN was performed on e11.5 peripheral blood (300K cells per replicate) using the EpiCypher CUTANA ChIC/CUT&RUN Kit. Replicates represent samples from a single embryo, with littermate controls for each genotype (Ncaph2 n=3, H4K20me1 in: Ncaph2 WT n=4, Ncaph2 Het n=3, KO n=4). Libraries were generated with the NEBNext Ultra II DNA Library Prep kit (NEB), optimized for small fragments. This included end preparation at 20°C for 30 minutes followed by 50°C for 60 minutes, AMPure XP cleanup at 1.75X for ligation, annealing at 65°C for library amplification, and a two-step AMPure XP cleanup (first at 0.8X negative selection, followed by 1.2X positive selection).

### Bioinformatics

RNA-seq reads were aligned to the mm10 reference genome using STAR, and quantified with featureCounts. Differential gene expression analysis was conducted using DESeq2 (Love et al., 2014)to compare *Ncaph2* Δ/Δ and Δ/+ conditions. Genes with an adjusted p-value < 0.05 were considered significant. Heatmaps of gene expression were generated using the pheatmap package in R, with Z-scores calculated from normalized expression values. Volcano plots displaying log2 fold change and adjusted p-values were created using ggplot in R. Gene ontology (GO) enrichment analysis for upregulated and downregulated genes was conducted using clusterProfiler, focusing on Biological Process (BP) terms with an adjusted p-value < 0.05.

CUT&RUN reads were aligned to the mm10 genome using Bowtie2, followed by duplicate removal with Picard. bamCoverage was used for RPKM normalization. Peaks were called using macs2 callpeak --broad for Ncaph2 and H3K27ac, and macs2 bdgpeakcall -c 200 for H4K20me1. Peak annotation was determined using ChIPseeker (Yu et al., 2015) annotatePeak, and Motif enrichment was determined using HOMER(Heinz et al., 2010) findMotifs.pl. Gene ontology (GO) enrichment analysis for peaks was conducted using clusterProfiler, focusing on Biological Process (BP) terms, and pathway enrichment was conducted using enrichPathway for genes associated with peaks (within 2kb of TSS). Heatmaps and profile plots of Ncaph2, H3K27ac, and H4K20me1 were generated in deepTools using computeMatrix and plotHeatmap or plotProfile over indicated regions. Peak intersection was calculated using bedtools intersect.

To compare Ncaph2 levels with RNA, H3K27ac, and H4K20me1, region-matched enrichment scores over gene promoters (TSS +/- 500 bp), gene bodies (TSS to TES), or merged exons (metagene option for RNA-seq) were generated in deepTools (Ramirez et al., 2014). For RNA, expressed transcripts were divided into quintiles based on Ncaph2 levels in R, and average normalized RNA was and plotted over the same quintiles. For H3K27ac and H4K20me1, all transcripts were divided into quintiles based on Ncaph2 levels in R, and average H3K27ac or H4K20me1 was plotted over the same quintiles. Density scatter plots were generated in R using ggplot and Pearson correlation coefficient calculated. Ncaph2 (TSS +/- 500bp) and H4K20me1 (TSS to TES) was quantified over differentially expressed genes using computeMatrix over custom bed file for upregulated, downregulated, and not significantly changed genes, and average scores were plotted over respective gene sets in R. Differential gene body H4K20me1 was determined using read count tables of H4K20me1 CUT&RUN average RPKM over gene bodies (TSS to TES) in KO vs. heterozygous samples as input for DESeq2.

To evaluate Ncaph2 occupancy across cell cycle phase transitions, genes sets for cell cycle phases (G1, S, G2, and M) were retrieved using enrichGO. computeMatrix was used to determine average Ncaph2 levels over promoters for each gene set. Heatmap and profile plot were generated in deepTools, and boxplot was generated in R.

Generation of Heatmap Demonstrating RNA Levels Over cell cycle-Related Genes: Custom bed files were used with computeMatrix to determine RNA over exons and Ncaph2 over promoters at indicated gene sets.

### Primitive Two-Step Culture Methods

Method adapted from (Malik et al., 2013). Briefly, E8.5 embryos from E8.5 *Ncaph2* FL/+ EpoR cre/+ x *Ncaph2* FL/FL crosses were dissected in PBS, washed, and transferred into individual tubes. For dissociation, 200 μL of 0.25% Trypsin was added to each tube, and the embryos were digested for 4 minutes with gentle titration every minute. Cells were then pelleted by centrifugation and resuspended in 500 μL of maturation medium, consisting of 1 mL PFHA, 1 mL serum replacement, 100 μL Glutamax, 7.8 mL 1X IMDM, 2 μL/mL Erythropoietin (Epo), and 1.28 μL of 1:10 MTG. Cells were seeded into 0.1% gelatin-coated 24-well plates, with one embryo per well, and incubated at 37°C in a humidified environment with high CO2. After 24 hours, 50 μL of additional maturation medium was added to each well. Cell counts were done daily. Viability was assessed via Trypan blue exclusion and benzidine staining was used to monitor cell growth and erythroid differentiation.

## References

Abbas, T., Shibata, E., Park, J., Jha, S., Karnani, N. & Dutta, A. 2010. CRL4(Cdt2) regulates cell proliferation and histone gene expression by targeting PR-Set7/Set8 for degradation. Mol Cell, 40, 9–21.

Abe, S., Nagasaka, K., Hirayama, Y., Kozuka-Hata, H., Oyama, M., Aoyagi, Y., Obuse, C. & Hirota, T. 2011. The initial phase of chromosome condensation requires Cdk1-mediated phosphorylation of the CAP-D3 subunit of condensin II. Genes Dev, 25, 863–74.

Bagger, F. O., Kinalis, S. & Rapin, N. 2019. BloodSpot: a database of healthy and malignant haematopoiesis updated with purified and single cell mRNA sequencing profiles. Nucleic Acids Res, 47, D881–D885.

Barski, A., Cuddapah, S., Cui, K., Roh, T. Y., Schones, D. E., Wang, Z., Wei, G., Chepelev, I. & Zhao, K. 2007. High-resolution profiling of histone methylations in the human genome. Cell, 129, 823–37.

Beck, D. B., Oda, H., Shen, S. S. & Reinberg, D. 2012. PR-Set7 and H4K20me1: at the crossroads of genome integrity, cell cycle, chromosome condensation, and transcription. Genes Dev, 26, 325–37.

Centore, R. C., Havens, C. G., Manning, A. L., Li, J. M., Flynn, R. L., Tse, A., Jin, J., Dyson, N. J., Walter, J. C. & Zou, L. 2010. CRL4(Cdt2)-mediated destruction of the histone methyltransferase Set8 prevents premature chromatin compaction in S phase. Mol Cell, 40, 22–33.

Dowen, J. M., Bilodeau, S., Orlando, D. A., Hubner, M. R., Abraham, B. J., Spector, D. L. & Young, R. A. 2013. Multiple structural maintenance of chromosome complexes at transcriptional regulatory elements. Stem Cell Reports, 1, 371–8.

Evertts, A. G., Manning, A. L., Wang, X., Dyson, N. J., Garcia, B. A. & Coller, H. A. 2013. H4K20 methylation regulates quiescence and chromatin compaction. Mol Biol Cell, 24, 3025–37.

Fox, S., Myers, J. A., Davidson, C., Getman, M., Kingsley, P. D., Frankiewicz, N. & Bulger, M. 2020. Hyperacetylated chromatin domains mark cell type-specific genes and suggest distinct modes of enhancer function. Nat Commun, 11, 4544.

Heinrich, A. C., Pelanda, R. & Klingmuller, U. 2004. A mouse model for visualization and conditional mutations in the erythroid lineage. Blood, 104, 659–66.

Heinz, S., Benner, C., Spann, N., Bertolino, E., Lin, Y. C., Laslo, P., Cheng, J. X., Murre, C., Singh, H. & Glass, C. K. 2010. Simple combinations of lineage-determining transcription factors prime cis-regulatory elements required for macrophage and B cell identities. Mol Cell, 38, 576–89.

Hirota, T., Gerlich, D., Koch, B., Ellenberg, J. & Peters, J. M. 2004. Distinct functions of condensin I and II in mitotic chromosome assembly. J Cell Sci, 117, 6435–45.

Hoencamp, C., Dudchenko, O., Elbatsh, A. M. O., Brahmachari, S., Raaijmakers, J. A., VAN Schaik, T., SEDENO Cacciatore, A., Contessoto, V. G., Van Heesbeen, R., Van Den Broek, B., Mhaskar, A. N., Teunissen, H., ST Hilaire, B. G., Weisz, D., Omer, A. D., Pham, M., Colaric, Z., Yang, Z., Rao, S. S. P., Mitra, N., Lui, C., Yao, W., Khan, R., Moroz, L. L., Kohn, A., ST Leger, J., Mena, A., Holcroft, K., Gambetta, M. C., Lim, F., Farley, E., Stein, N., Haddad, A., Chauss, D., Mutlu, A. S., Wang, M. C., Young, N. D., Hildebrandt, E., Cheng, H. H., Knight, C. J., Burnham, T. L. U., Hovel, K. A., Beel, A. J., Mattei, P. J., Kornberg, R. D., Warren, W. C., Cary, G., GOMEZ-Skarmeta, J. L., Hinman, V., Lindblad-Toh, K., DI Palma, F., Maeshima, K., Multani, A. S., Pathak, S., NEL-Themaat, L., Behringer, R. R., Kaur, P., Medema, R. H., VAN Steensel, B., DE Wit, E., Onuchic, J. N., DI Pierro, M., Lieberman Aiden, E. & Rowland, B. D. 2021. 3D genomics across the tree of life reveals condensin II as a determinant of architecture type. Science, 372, 984–989.

Ji, P., Murata-Hori, M. & Lodish, H. F. 2011. Formation of mammalian erythrocytes: chromatin condensation and enucleation. Trends Cell Biol, 21, 409–15.

Jorgensen, S., Schotta, G. & Sorensen, C. S. 2013. Histone H4 lysine 20 methylation: key player in epigenetic regulation of genomic integrity. Nucleic Acids Res, 41, 2797–806.

Kingsley, P. D., Malik, J., Fantauzzo, K. A. & Palis, J. 2004. Yolk sac-derived primitive erythroblasts enucleate during mammalian embryogenesis. Blood, 104, 19–25.

Koulnis, M., Pop, R., Porpiglia, E., Shearstone, J. R., Hidalgo, D. & Socolovsky, M. 2011. Identification and analysis of mouse erythroid progenitors using the CD71/TER119 flow-cytometric assay. J Vis Exp.

Lancaster, L., Patel, H., Kelly, G. & Uhlmann, F. 2021. A role for condensin in mediating transcriptional adaptation to environmental stimuli. Life Sci Alliance, 4.

Liu, W., Tanasa, B., Tyurina, O. V., Zhou, T. Y., Gassmann, R., Liu, W. T., Ohgi, K. A., Benner, C., Garcia-Bassets, I., Aggarwal, A. K., Desai, A., Dorrestein, P. C., Glass, C. K. & Rosenfeld, M. G. 2010. PHF8 mediates histone H4 lysine 20 demethylation events involved in cell cycle progression. Nature, 466, 508–12.

Love, M. I., Huber, W. & Anders, S. 2014. Moderated estimation of fold change and dispersion for RNA-seq data with DESeq2. Genome Biol, 15, 550.

Macdonald, L., Taylor, G. C., Brisbane, J. M., Christodoulou, E., Scott, L., VON Kriegsheim, A., Rossant, J., Gu, B. & Wood, A. J. 2022. Rapid and specific degradation of endogenous proteins in mouse models using auxin-inducible degrons. Elife, 11.

Malik, J., Getman, M. & Steiner, L. A. 2015. Histone methyltransferase Setd8 represses Gata2 expression and regulates erythroid maturation. Mol Cell Biol, 35, 2059–72.

Malik, J., Kim, A. R., Tyre, K. A., Cherukuri, A. R. & Palis, J. 2013. Erythropoietin critically regulates the terminal maturation of murine and human primitive erythroblasts. Haematologica, 98, 1778–87.

Malik, J., Lillis, J. A., Couch, T., Getman, M. & Steiner, L. A. 2017. The Methyltransferase Setd8 Is Essential for Erythroblast Survival and Maturation. Cell Rep, 21, 2376–2383.

Myers, J. A., Couch, T., Murphy, Z., Malik, J., Getman, M. & Steiner, L. A. 2020. The histone methyltransferase Setd8 alters the chromatin landscape and regulates the expression of key transcription factors during erythroid differentiation. Epigenetics Chromatin, 13, 16.

Nishide, K. & Hirano, T. 2014. Overlapping and non-overlapping functions of condensins I and II in neural stem cell divisions. PLoS Genet, 10, e1004847.

Oda, H., Hubner, M. R., Beck, D. B., Vermeulen, M., Hurwitz, J., Spector, D. L. & Reinberg, D. 2010. Regulation of the histone H4 monomethylase PR-Set7 by CRL4(Cdt2)-mediated PCNA-dependent degradation during DNA damage. Mol Cell, 40, 364–76.

Oda, H., Okamoto, I., Murphy, N., Chu, J., Price, S. M., Shen, M. M., Torres-Padilla, M. E., Heard, E. & Reinberg, D. 2009. Monomethylation of histone H4-lysine 20 is involved in chromosome structure and stability and is essential for mouse development. Mol Cell Biol, 29, 2278–95.

Ono, T., Fang, Y., Spector, D. L. & Hirano, T. 2004. Spatial and temporal regulation of Condensins I and II in mitotic chromosome assembly in human cells. Mol Biol Cell, 15, 3296–308.

Ono, T., Sakamoto, C., Nakao, M., Saitoh, N. & Hirano, T. 2017. Condensin II plays an essential role in reversible assembly of mitotic chromosomes in situ. Mol Biol Cell, 28, 2875–2886.

Palis, J., Malik, J., Mcgrath, K. E. & Kingsley, P. D. 2010. Primitive erythropoiesis in the mammalian embryo. Int J Dev Biol, 54, 1011–8.

Ramirez, F., Dundar, F., Diehl, S., Gruning, B. A. & Manke, T. 2014. deepTools: a flexible platform for exploring deep-sequencing data. Nucleic Acids Res, 42, W187–91.

Shoaib, M., Chen, Q., Shi, X., Nair, N., Prasanna, C., Yang, R., Walter, D., Frederiksen, K. S., Einarsson, H., Svensson, J. P., Liu, C. F., Ekwall, K., Lerdrup, M., Nordenskiold, L. & Sorensen, C. S. 2021. Histone H4 lysine 20 mono-methylation directly facilitates chromatin openness and promotes transcription of housekeeping genes. Nat Commun, 12, 4800.

Skene, P. J. & Henikoff, S. 2017. An efficient targeted nuclease strategy for high-resolution mapping of DNA binding sites. Elife, 6.

Woodward, J., Taylor, G. C., Soares, D. C., Boyle, S., Sie, D., Read, D., Chathoth, K., Vukovic, M., Tarrats, N., Jamieson, D., Campbell, K. J., Blyth, K., Acosta, J. C., Ylstra, B., Arends, M. J., Kranc, K. R., Jackson, A. P., Bickmore, W. A. & Wood, A. J. 2016. Condensin II mutation causes T-cell lymphoma through tissue-specific genome instability. Genes Dev, 30, 2173–2186.

Xu, Y., Leung, C. G., Lee, D. C., Kennedy, B. K. & Crispino, J. D. 2006. Mtb, the murine homolog of condensin II subunit CAP-G2, represses transcription and promotes erythroid cell differentiation. Leukemia, 20, 1261–9.

Yu, G., Wang, L. G. & He, Q. Y. 2015. ChIPseeker: an R/Bioconductor package for ChIP peak annotation, comparison and visualization. Bioinformatics, 31, 2382–3.

Yuen, K. C., Slaughter, B. D. & Gerton, J. L. 2017. Condensin II is anchored by TFIIIC and H3K4me3 in the mammalian genome and supports the expression of active dense gene clusters. Sci Adv, 3, e1700191.

